# Robust odor coding across states in piriform cortex requires recurrent circuitry: evidence for pattern completion in an associative network

**DOI:** 10.1101/694331

**Authors:** Kevin A. Bolding, Shivathmihai Nagappan, Bao-Xia Han, Fan Wang, Kevin M. Franks

**Author notes:** Correspondence: Kevin Franks, Bryan Research Building, Rm. 401D, 311 Research Dr., Durham, NC, 27705. Author contributions: K.A.B. and K.M.F. conceived the experiments, analyzed the experiments and wrote the paper. K.A.B. performed the experiments. B.H. and F.W. made the Ntng1-cre mouse line. S.N. validated the Ntng1-cre line and S.N. performed experiments on Ntng1-cre mice with K.A.B.

## Abstract

Pattern completion, or the ability to retrieve stable neural activity patterns from noisy or partial cues, is a fundamental feature of memory. Theoretical studies indicate that recurrently connected auto-associative or discrete attractor network models can perform this process. Although phenomenological evidence for pattern completion and attractor dynamics have been described in various recurrent neural circuits, the crucial role that recurrent circuitry plays in implementing these processes has not been shown. Here we show that although odor representations in mouse olfactory bulb degrade under anesthesia, responses in downstream piriform cortex remain robust. Recurrent connections are required to stabilize cortical odor representations across states. Moreover, piriform odor representations exhibit attractor dynamics, both within and across trials, and these are also abolished when recurrent circuitry is eliminated. Thus, an auto-associative cortical circuit stabilizes output in response to degraded input, and the recurrent circuitry that defines these networks is required for this stabilization.

## Main

Recognition occurs at the interface of perception and memory: it requires being able to reliably identify a familiar object, even when the stimulus is noisy or degraded, or when behavioral states change. Sensory systems must therefore be able to generate representations of the stimuli that are robust to changes in input or ongoing brain activity. Theoretical studies have shown that this retrieval can occur in recurrent neural networks through processes called auto-associative recall, attractor dynamics or pattern completion^1-9^. Phenomenological evidence for pattern completion has been described in various recurrent networks, including hippocampus^10-12^, piriform cortex^13^, and neocortex^14-16^. Despite recurrent collateral connections being a defining feature of auto-associative networks, whether these phenomena indeed require recurrent connectivity remains unclear.

Odors are initially encoded as combinations of co-active glomeruli in the olfactory bulb (OB). This elemental odor code is then integrated in piriform cortex (PCx) to form a synthetic or gestalt odor representation of the input set. Projections from OB to PCx are diffuse and overlapping^17-19^ allowing individual principal neurons in PCx to integrate inputs from different and possibly random subsets of glomeruli. The diffuse projections from OB to PCx activate odor-specific ensembles of neurons distributed across PCx^20,21^ whose concerted activity encodes odor identity ^22,23^. Principal cells make excitatory synaptic connections onto other PCx cells with sparse but uniform connection probabilities across millimeters of PCx^24,25^, forming an extensive recurrent network, similar in synaptic organization to hippocampal CA3^8^. Theoretical studies have shown how such recurrently connected ensembles can generate stable odor representations that are robust to partial input^4,26,27^. Thus, PCx is thought to resemble an auto-associative network that can form stable cortical odor representations. Whether recurrent circuits are actually required to stabilize output patterns given noisy or partial input has not been shown.

PCx populations can demonstrate pattern separation or pattern completion-like responses depending on recent training history^13^. However, these and other major observations regarding associative functions and odor-evoked activation patterns in PCx were obtained in either ex vivo^28,29^ or anesthetized preparations^20,30^. Odor responses in both OB and PCx are reported to depend strongly on global brain state^31-33^. We set out to determine how responses differ in PCx in awake and anesthetized conditions. We found degraded OB responses but little difference in the quality of PCx representations suggesting that PCx circuits may transform inputs to stabilize odor representations across conditions. Using a novel transgenic line and selective disruption of PCx output, we localized this function to the recurrent connections within PCx.

## Results

### Odor responsivity is state-dependent in OB but not PCx

We simultaneously recorded spiking activity in populations of mitral cells in OB and layer II neurons in PCx in head-fixed mice before and after inducing ketamine/xylazine anesthesia (k/x; Fig. 1a). Anesthesia induced pronounced changes in respiration patterns, oscillatory activity and spontaneous spiking in both OB and PCx (Fig. 1, b and c; Supplementary Fig. 1), consistent with previous studies^31,33-35^. We examined odor-evoked spiking activity in individual cells in OB and PCx during the first sniff after odor delivery (Fig. 1, d and e). OB neurons were less responsive under anesthesia, with many fewer significantly activated neurons and increased lifetime and population sparseness (Fig. 1, f and g). We took care to ensure that this effect was indeed due to changes in spiking activity and not to unit drift or instabilities over the course of the recording (Supplementary Fig. 2). Yet, despite reduced input from OB, PCx responsivity shifted only subtly and, in fact, toward greater activation (Figs. 1, h and i; Supplementary Fig. 3), due largely to an increase in signal-to-noise ratio as spontaneous activity levels in PCx dropped under anesthesia.

**Figure 1.**
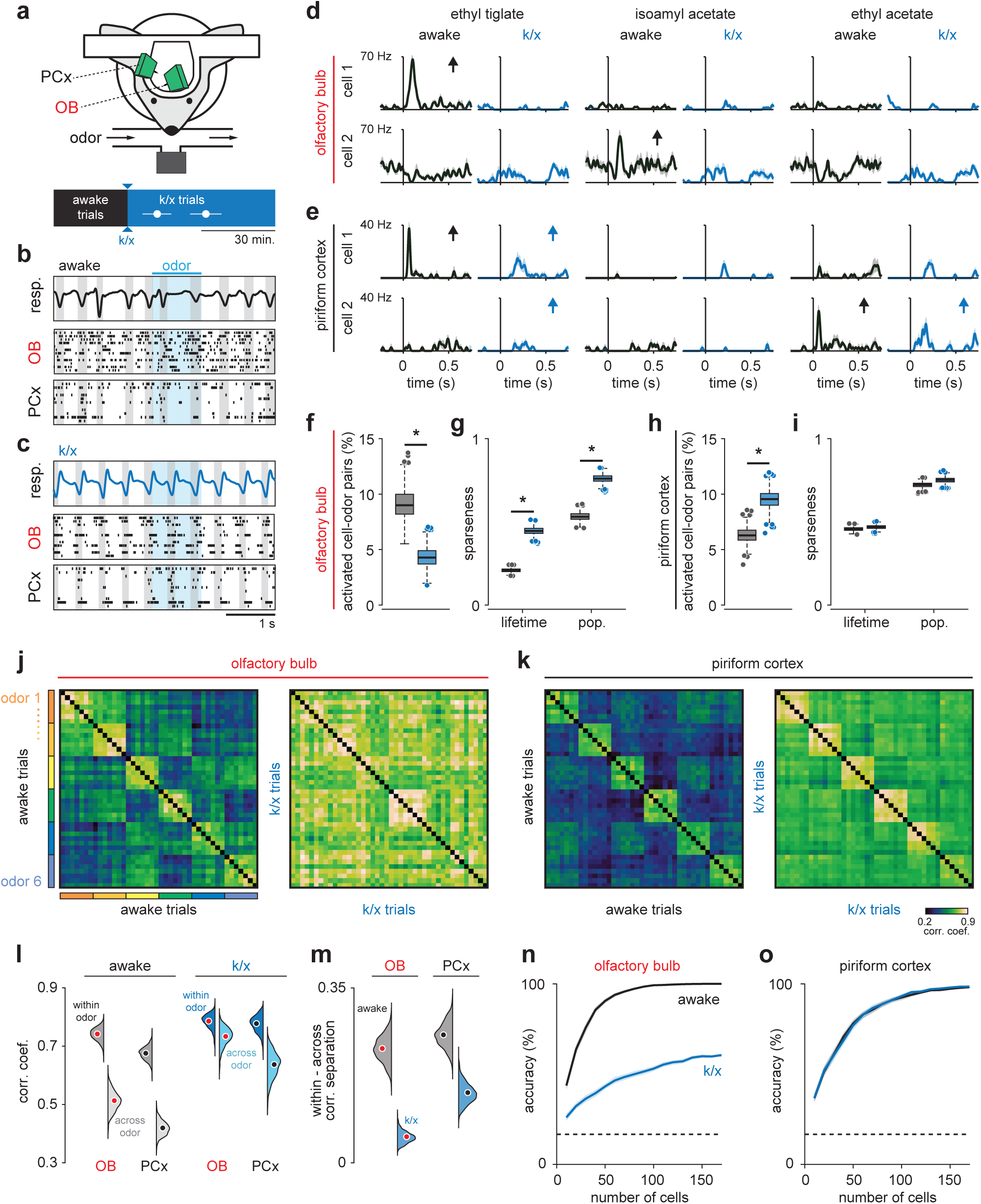
Odor responsivity is state-dependent in OB but not PCx. (a) Recording and experimental schematics. Timing of first and last trials designated as anesthetized for analysis, starting from the onset of behavioral indicators of anesthesia and continuing for 10-15 presentations of all odors (n = 12 experiments in 6 mice). Error bars are mean ± SEM. (b) Example recording showing irregular respiration (top) and desynchronized spiking in awake OB (middle) and PCx (bottom). Ticks indicate action potentials and each row represents a different cell. Gray shading indicates inhalation, blue shading indicates odor delivery. (c) Same recording as b after k/x/ injection. Respiration becomes rhythmic and spontaneous spiking in OB and PCx slows and becomes entrained to respiration under anesthesia. (d) Example OB cell-odor pair responses during awake and k/x trials showing loss of significant odor responses under anesthesia. Arrows indicate significant responses. (e) Example PCx cell-odor pair responses during awake and k/x trials showing preserved odor responses under anesthesia. Arrows indicate significant responses. (f) Percent of OB cell-odor pairs with significant activation (n = 187 cells, 12 experiments). Asterisks indicate p < 0.05 in bootstrap difference test (p = 0.007). Boxes indicate quartiles and whiskers indicate ± 2.7 st. dev. from mean. Data points outside this range are shown as circles. (g) Lifetime and population sparseness in OB (bootstrap difference test, p < 0.05) across all cells and odors. Asterisks indicate p < 0.05 on bootstrap difference test. Lifetime: p < 0.001; Population: p < 0.001. (h-i) As in f-g, but in PCx (n = 640 cells, 11 experiments). Activation: p < 0.001; Lifetime: p = 0.45; Population: p = 0.22. (j-k) Trial-by-trial pseudopopulation correlation matrices sorted by odor within OB (j) and PCx (k) in awake and anesthetized conditions. (l) Bootstrapped distributions of withinand across-odor trial-to-trial correlations in OB and PCx computed by sampling cells with replacement and computing the mean correlation within and across odors 1000 times. Means are shown as filled circles. (m) Separation (mean within-odor correlations – mean across-odor correlations) of odor representations in OB and PCx over 1000 bootstrap iterations. (n-o) Odor classification accuracy as a function of pseudopopulation size in OB (n) and PCx (o) in awake and anesthetized states using a multiclass linear support vector machine. Mean ± 95% bootstrapped confidence intervals.

To examine odor responses at the population level we constructed pseudopopulation vectors of OB or PCx firing rates for each odor trial and measured the similarity of responses using trial-to-trial correlations within or across odors (Fig. 1, j-l). In these and other analyses, we excluded the first ∼5 trials to minimize the contribution of rapid sniffing in response to the first few odor presentations, and we used an equivalent number of trials during stable anesthesia, after initial induction but before signs of recovery (Fig. 1a). Although responses to different odors became more correlated under anesthesia in both regions, this difference was substantially more pronounced in OB than PCx (Fig. 1m). Greater inter-odor response separation in PCx than OB under anesthesia was also observed when comparing population vector correlations for simultaneously recorded OB and PCx neurons (Supplementary Fig. 4). We next determined how accurately odors could be identified from spiking activity of populations of OB and PCx neurons. To do this, we trained and tested classifiers on either awake or anesthetized odor trials from pseudopopulations to match population sizes between OB or PCx neurons. Classifier performance using individual PCx experiments with simultaneously recorded populations of up to 60 cells did not differ substantially from accuracy for pseudopopulations of the same size (Supplementary Fig. 4). Decoding accuracy was dramatically worse under anesthesia in OB (Fig. 1n), whereas responses in PCx were decoded equally well in either state (Fig. 1o).

### PCx stabilizes odor representations across activity regimes

These data indicate that PCx maintains odor responsivity when OB input is degraded, but not whether the cortical odor representations themselves are preserved across states. To address this, we focused first on cells that had significant increases in odor-evoked spiking. We defined activated cell-odor pairs as either *robust* if they were activated in both regimes or as *state-specific* if they were only activated in awake or only in anesthetized states (Fig. 2, a and b). There were many more *robust* responses in PCx than OB (OB, 21/102 total awake responses; PCx, 118/237 total awake responses, Fig. 2c). To avoid simply classifying responses as ‘activated’ or ‘not-activated’ according to an arbitrary statistical threshold, we again considered single-trial population responses using spike rates for all OB and PCx neurons. Compared to OB, PCx population responses to the same odor were more correlated across states (Fig. 2d) and were more separable from responses to distinct odors (Fig. 2e), indicating that population odor representations are better preserved across states in PCx. To determine whether cross-state response correlations reflect stable odor representations, we trained a classifier using responses recorded in the awake state and tested the classifier on responses recorded under anesthesia. Cross-state decoding was substantially better in PCx than OB (Fig. 2f), indicating that PCx extracts and selectively represents stimulus-specific information from partial and noisy OB input. Using Principal Components Analysis (PCA) we found that state accounted for most of the variance in responses across states, in both regions (Supplementary Fig. 5). If pattern stability is a reflection of overlap in a low-dimensional neural activity space, then neither region maintained similar responses across states. We considered an alternative model in which a downstream decoder could adapt to overall state-dependent changes and still maintain the ability to distinguish stimuli. To explore this possibility, we used demixed PCA to isolate the stimulus-specific features of the population responses ^36^. This analysis revealed that OB responses to different odors were clearly separable and overlapped partly in awake and anesthetized states (Fig. 2g). However, responses overlapped almost completely in PCx, indicating that the stimulus-specific features of the cortical odor response are almost identical across states and better preserved than their inputs (Fig. 2, h and i). Thus, regardless of whether downstream processing is fixed or adaptive, PCx actively transforms degraded OB output to represent the stimulus more faithfully than the input it receives.

**Figure 2.**
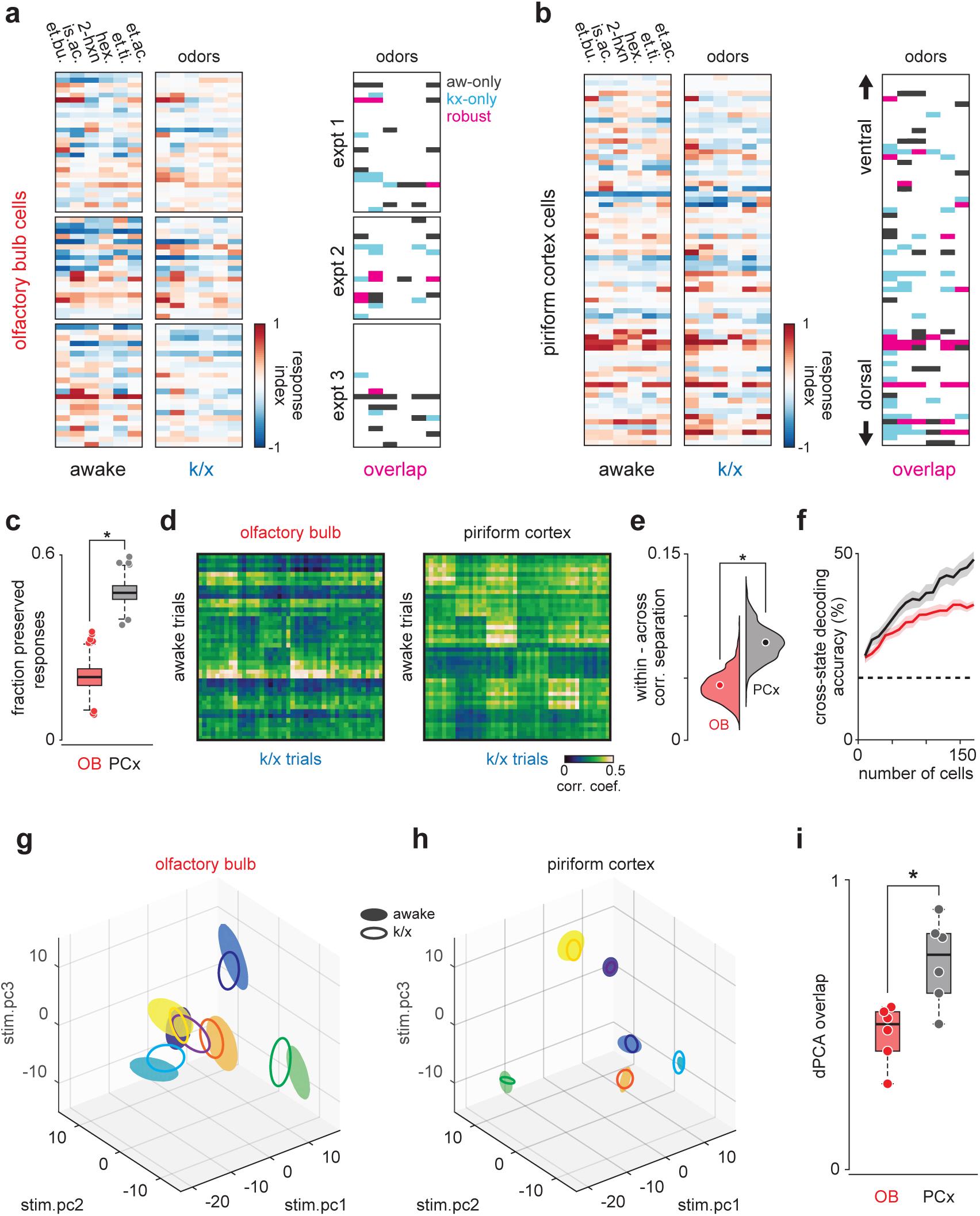
PCx stabilizes odor representations across activity regimes. (a-b) Example population responses to six odors for three OB (a) and one PCx recordings (b). Awake-only (black), k/x-only (cyan), and robust cell-odor pairs, which were activated in both states (magenta, p < 0.05 rank-sum test), are indicated at right. PCx cells are sorted by dorsal-ventral location. Note more robust PCx responses in deeper, (i.e. more dorsal) layer II. (c) Fraction of significant awake cell-odor pair responses that are preserved under anesthesia in OB (n = 187 cells, 12 experiments) and PCx (n = 640 cells, 11 experiments). Asterisks indicate p < 0.05 on bootstrap difference test (p < 0.001). (d) Cross-state trial-by-trial pseudopopulation correlation matrices sorted by odor within OB and PCx illustrate similarity between awake and anesthetized population responses. (e) Separation (mean within-odor correlations – mean across-odor correlations) of cross-state odor representations in OB and PCx over 1000 bootstrap iterations. Asterisks indicate p < 0.05 in bootstrap difference test (OB vs PCx, p < 0.001). Means are shown as filled circles. (f) Cross-state decoding accuracy (trained on awake trials, tested on k/x trials) in OB and PCx. Mean ± 95% bootstrapped confidence intervals. (g) OB pseudopopulation activity projected onto the first three stimulus-dependent demixed PCA components and shown as mean ± 1 s.d. ellipsoids across trials of the same odor. Different colors correspond to different odors. Filled ellipsoids are awake, unfilled ellipsoids are anesthetized. (h) As in g, but for PCx. Note overall larger separation between odors in either state. (i) Overlap between awake and k/x trial response distributions projected onto first three stimulus-dependent components in OB (red) and PCx (black). Dots show overlap scores for individual odors. Overlap is calculated using Matusita’s overlap measure (see Methods). Unpaired t-test, n = 6 odors, t(10) = −3.41, p = 0.007.

### Robust PCx representations derive from short-latency OB responses

To generate stable cortical odor representations using degraded and noisy OB input, PCx must over-represent the impact of the few *robust* OB responses. What features of the OB response does PCx use to selectively extract this information? Peak firing rate distributions of *robust* and *state-specific* OB responses were broad and overlapped substantially in either regime (Fig. 3, a and c), and though robust responses appeared slightly larger on average this difference was not statistically significant. However, *robust* OB responses had significantly shorter latencies than *state-specific* responses (Fig. 3, a and d). Thus, PCx could over-represent early inputs to produce a stable output^37^. Indeed, *robust* PCx responses had shorter latencies (Fig. 3, b and f), but *robust* PCx responses were also substantially stronger than *state-specific* ones (Fig. 3, b and e), indicating that *robust* responses are actively amplified within PCx. Notably, theoretical models have shown that responses in a recurrently-connected network can track input changes nearly-instantaneously^38^, whereas feedforward networks respond at a temporal delay, suggesting that recurrent excitation within PCx could facilitate short-latency, robust responses. PCx also sends centrifugal inputs to OB, and could return an accurate representation of the current stimulus to the OB, allowing comparison between ongoing input and an internally-generated prediction based on the retrieved activation pattern^39^. This process may augment the selective representation of early OB inputs in PCx.

**Figure 3.**
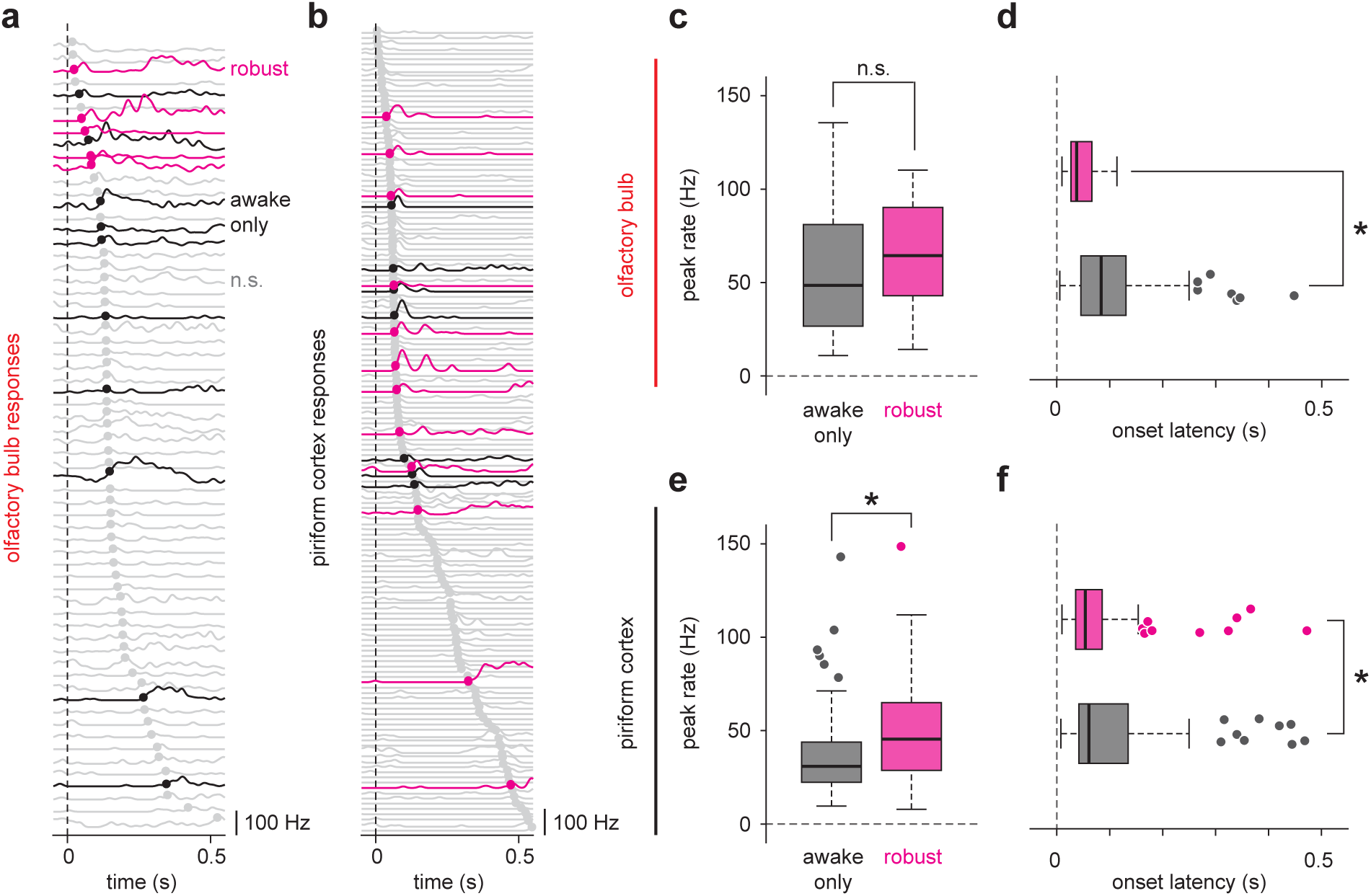
Robust PCx representations derive from short-latency OB responses. (a-b) All cell-odor pair responses for example simultaneously recorded OB (a) and PCx (b) populations sorted by onset latency determined by 2.5 st. dev. threshold crossing. Latencies are marked with filled circles. Robust, awake-only, and non-significant responses are magenta, black, and gray, respectively. (c-d) Peak firing rates (c) and onset latencies (d) for robust (magenta) vs awake-only (black) responses in OB. Boxes indicate quartiles and whiskers indicate ± 2.7 st. dev. from mean. Data points outside this range are shown as circles. Asterisks indicate p < 0.05 in unpaired t-test. n = 81 awake-only OB cell-odor pairs and 21 robust OB cell-odor pairs. Peak: t(100) = −0.82, p = 0.42; Latency: t(100) = 3.06, p = 0.003. (e-f) As in c-d but for PCx. n = 125 awake-only PCx cells and 116 robust PCx cells. Peak: t(239) = −4.49, p = 1.12e-5; Latency: t(236) = 2.47, p = 0.01.

### Pattern recovery depends on PCx recurrent circuits

Piriform cortex is a recurrent cortical circuit that resembles auto-associative or discrete attractor networks. The ability to generate stable output patterns using degraded input is a property of such networks. If PCx is indeed such a network then the recurrent connectivity between PCx neurons should be essential for stabilizing odor representations across states. PCx contains two distinct classes of principal cells: pyramidal cells (PYRs), which are located primarily in deeper layer II and receive both OB and recurrent collateral inputs, and semilunar cells (SLs), which are more superficial and receive strong OB input but no recurrent input (Fig. 4a)^40,41^. To test whether recurrent connectivity predicts response stability, we generated a Netrin G1-Cre mouse line (*Ntng1*-Cre) that selectively expresses Cre recombinase in SLs in PCx (Fig. 4, b and c). We then virally expressed either Cre-dependent Archaeorhodopsin-3 (Arch), Jaws, or Channelrhodopsin-2 (ChR2) in PCx to optogenetically differentiate SLs from PYRs during population recordings (Fig. 4, d-f, Supplementary Fig. 6). Indeed, odor responses were substantially more well-preserved across states in PYRs than SLs: (Fig. 4, g and h). Furthermore, SLs recapitulated the degradation of representations under anesthesia (Fig. 4, i and j) and the poor cross-state decoding (Fig. 4k) observed in OB output, whereas PYRs successfully recovered representations across states. Thus, two PCx cell populations, receiving the same feedforward input but distinguished by their recurrent connectivity, differentially recover odor representations when OB input is degraded.

**Figure 4.**
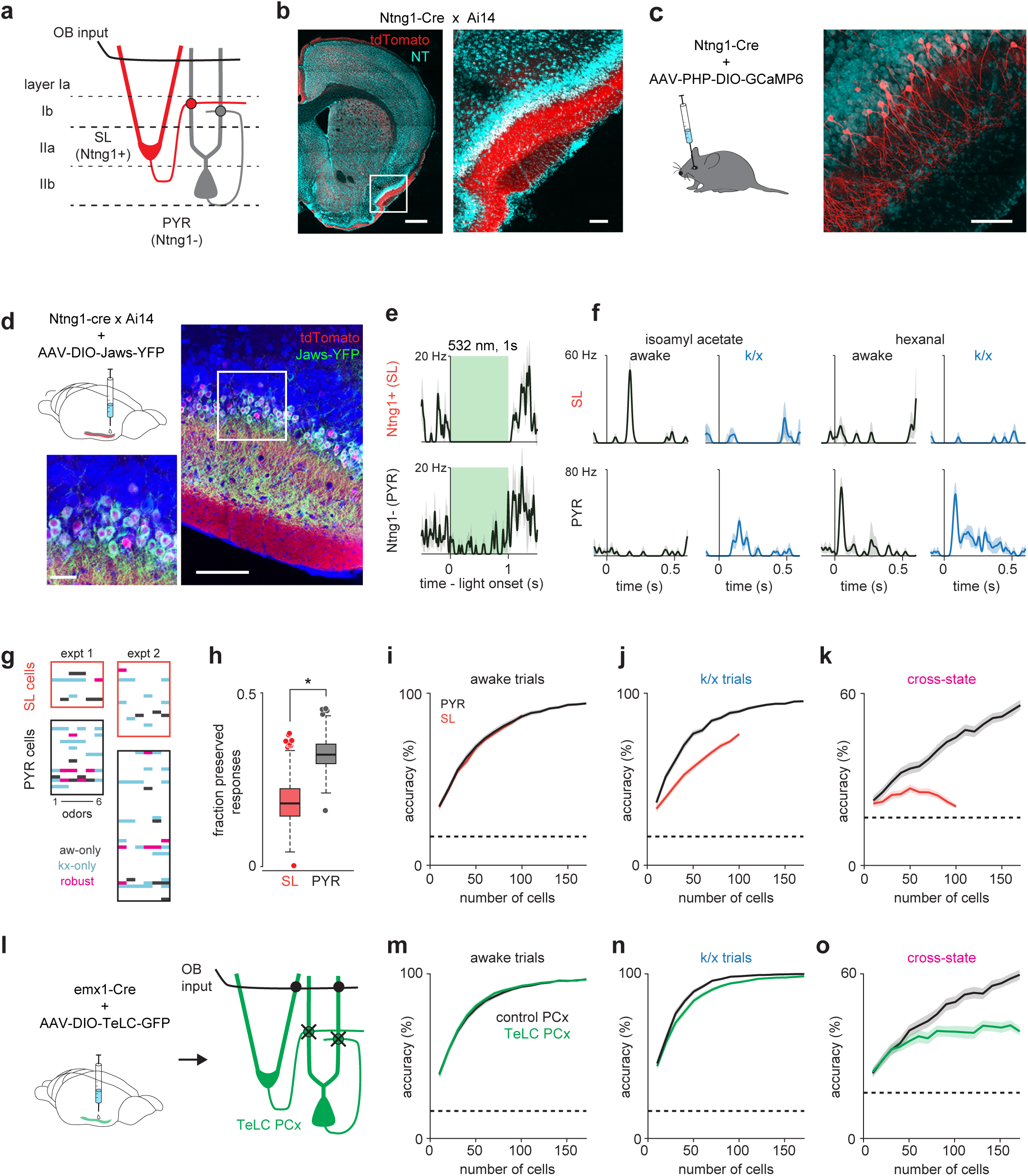
Pattern recovery depends on PCx recurrent circuits. (a) Schematic of inputs to excitatory cell-types in PCx. Semilunar cells receive OB input; pyramidal cells receive OB and recurrent collateral inputs. (b-c) Selective expression in PCx semilunar cells using Ntng1-Cre driver line. (b) Strong expression of Cre-dependent tdTomato in PCx layer II of Ntng1-Cre X Ai14 mice. Scale bars: 500 μm and 100 μm. (c) Sparse Cre-dependent GCaMP6 expression shows Ntng1+ cells restricted to superficial layer II and lacking basal dendrites. Scale bar: 100 μm. (d) Identifying Ntng1+ (semilunar, SL) and Ntng1-(pyramidal, PYR) cells in vivo using optogenetic inhibition. Injection of AAV expressing Cre-dependent Jaws in anterior piriform cortex produces selective expression in cells with semilunar localization and morphology. Scale bars: 100 μm and 20 μm. (e) Simultaneously recorded example cells exhibiting suppression (top) or residual spiking (bottom) in response to 1s, 532 nm light pulses. Mean ± SEM responses over 40 laser pulses. Optogenetic tagging and unit stability criteria identified 108 SL and 234 PYR cells in 12 experiments from 7 mice. (f) Odor responses during awake and k/x trials for an example SL (top) and PYR cell (bottom). (g) State-specific (black, awake-only; cyan k/x only) and robust (magenta) responses in simultaneously recorded populations of SL and PYR cells from two example experiments. (h) Fraction of significant awake cell-odor pair responses that are preserved under anesthesia in SL and PYR. Asterisk indicates p < 0.05 on bootstrap difference test (p = 0.036). (i-j) Odor classification accuracy as a function of pseudopopulation size using SL (red) and PYR (black) cells in awake (H) and anesthetized (I) states. Mean ± 95% bootstrapped confidence intervals. (k) Cross-state decoding accuracy using SL (red) and PYR (black) cells. Mean ± 95% bootstrapped confidence intervals. (l) Strategy for disabling recurrent circuits in PCx. Expression of tetanus toxin in all PCx excitatory cells disrupts recurrent connectivity. (m-n) Odor classification accuracy as a function of pseudopopulation size in TeLC-infected (green) and contralateral control (black) PCx in awake (m) and anesthetized (n) states. Mean ± 95% bootstrapped confidence intervals. Pseudopopulations were built from 241 cells recorded in control hemisphere in 4 experiments with 4 mice and 214 cells from TeLC hemisphere in 6 experiments with 5 mice. (o) Cross-state decoding accuracy in TeLC- (green) and control (black) PCx. Mean ± 95% bootstrapped confidence intervals.

Next, we asked whether eliminating recurrent connectivity in PCx abolished the ability to recover odor representations across states. To test this prediction, we injected conditional viral vectors to express tetanus toxin (AAV-DIO-GFP-TeLC) into PCx of *emx1*-Cre mice to express TeLC in all PCx excitatory neurons (Fig. 4l), effectively converting PCx into a pure feedforward circuit driven by OB^37^. We then obtained simultaneous bilateral recordings from TeLC-infected and contralateral control PCx hemispheres before and during anesthesia. Odor responses could be classified accurately in awake responses from both control and TeLC PCx (Fig. 4m). Somewhat surprisingly, TeLC-PCx decoding under anesthesia was not impaired (Fig. 4n). Because TeLC expression interferes with all PCx output, including strong feedback projections from PCx to OB inhibitory neurons, OB responses ipsilateral to TeLC-PCx can be substantially enhanced^37^, and may no longer present degraded input to PCx under anesthesia. Indeed, awake and anesthetized responses in OB ipsilateral to TeLC-PCx were decoded with equivalent accuracy, whereas simultaneous recordings from contralateral OB degraded under anesthesia (Supplementary Fig. 7). Thus, under anesthesia, PCx receives less accurate odor responses from the control OB than from TeLC-ipsilateral OB, but produces equivalent output, suggesting that recurrent circuitry accommodates worse OB input to improve decoding within PCx. Control PCx responses also provided for accurate cross-state decoding (Fig. 4o), whereas decoding from control OB responses was degraded (Supplementary Fig. 7), replicating our original observations from simultaneous OB-PCx recordings (Fig. 2f). Critically, cross-state decoding in PCx was markedly degraded in TeLC PCx (Fig. 4o), indicating that recovery of the odor-specific features of the cortical response during anesthesia requires recurrent connections.

### Rapid pattern formation in PCx populations

The enhanced stability of pattern activation across conditions in PCx is consistent with a pattern completion-like process occurring in PCx recurrent circuits. For stable population activity patterns to be retrieved they must first be formed, and this is commonly thought to occur through repeated experience with specific stimuli. We therefore sought evidence for odor pattern formation in PCx through experience over early trials in our recordings. If learning occurs during these early trials then responses should systematically converge toward a stable pattern. We tested this prediction by measuring distances in neural activity space for population responses. Responses on initial trials were highly variable but then stabilized over subsequent trials (Fig. 5, a-c). Much of this variability can be explained by increased sniffing (Fig. 5d), and therefore stronger overall responses, on early trials, however a multiple linear regression analysis revealed a significant effect of trial number alone (Fig. 5e). Recording instabilities or changes in cell health did not account for this shift as corresponding changes did not occur for pre-trial activity (Fig. 5b). We considered that some part of this increased stability could be inherited from and imparted to OB. Though our ability to interpret single-trial OB responses is limited by relatively low simultaneously recorded population sizes, we found OB responses indeed stabilized over early trials (Fig. 5f-j). OB stabilization dynamics were more rapid that in PCx and there was a specific decrease in odor-evoked overall firing rates in PCx that was not apparent in OB. However, the contribution of centrifugal inputs from PCx to OB make it impossible to determine whether the trial-dependent effects observed in OB reflects changes in OB that are then imparted to PCx or changes in PCx that are then imparted to OB.

**Figure 5.**
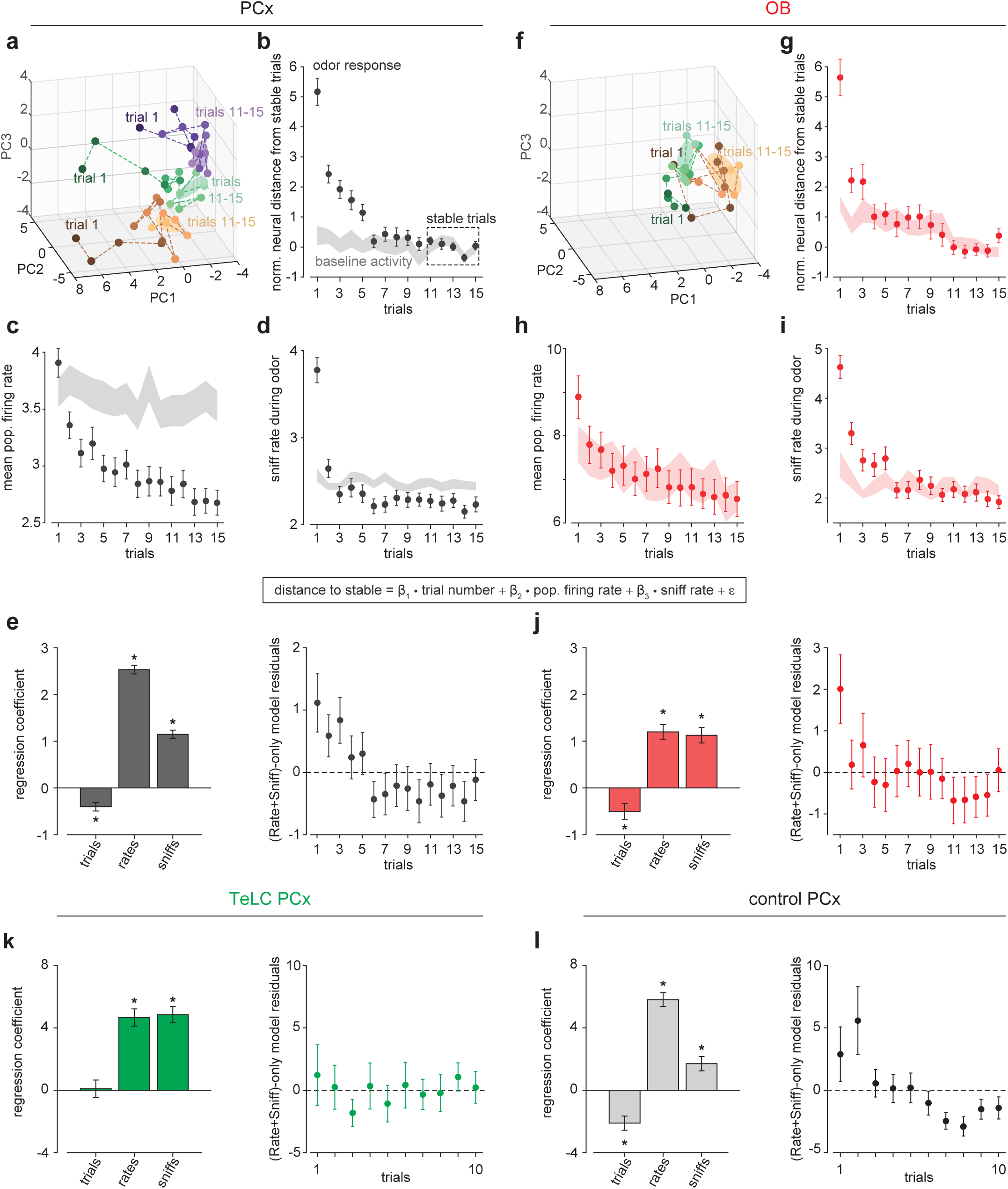
Rapid pattern formation in PCx population responses. (a, f) PCA trajectories for the first 15 presentations of three odors in an example simultaneous PCx (a) and OB (f) recording. The third odor in OB data occupied overlapping PC space and was omitted for visual clarity. The area occupied during the designated ‘stable’ trials are shown as mean ± 1 s.d. ellipsoids. Different colors correspond to different odors. (b, g) Average Euclidean distance from trial population vectors to stable trials normalized by the average distance between stable trials (b, PCx: n = 132 experiment-odor pairs, g, OB: n = 45 experiment-odor pairs, mean ± SEM). Shaded area shows distances computed using pre-odor baseline activity (mean ± SEM). (c, h) Average population firing rates during odor response (mean ± SEM) or pre-odor baseline (shaded area, mean ± SEM) for PCx (c) or OB (h). (d, i) Average sniffing rates during odor response (mean ± SEM) or pre-odor baseline (shaded area, mean ± SEM) for PCx recordings (d) and OB recordings (i). (e) Left, multiple linear regression coefficients for effects of sniff rate, population firing rate, and trial number on population distance to stable in PCx (mean ± SE). All main and interaction coefficients are significant (p < 0.05). Right, Residuals plot of multiple linear regression on distance-to-stable fit with only sniff rate and population firing rate predictors, showing decreasing distance with trial number independent of these predictors. (j) As in e, but for OB recordings. (k, l) As in e, but for TeLC-infected PCx (k, n = 36 experiment-odor pairs) and contralateral control PCx (l, n = 24 experiment-odor pairs). Distance changes are fully explained by sniff rate and overall population firing rate in TeLC-PCx, but depend on trial number in control PCx.

To account for the effects of receptor desensitization, sniffing, arousal, and the confounding contribution of centrifugal inputs, we compared the stabilization of responses across trials in bilateral recordings from control versus TeLC-infected PCx in the same animals. Under these conditions, we again observed a significant trial-number effect in contralateral control hemispheres, but sniffing and overall rate could fully account for response variability when we performed the same analysis on TeLC-infected PCx, with no significant trial number effect (Fig. 5, k and l). These data support the hypothesis that the trial number effect represents learning instantiated in PCx recurrent circuitry.

### Short-term pattern stability in recurrently-connected PCx cells

Activity patterns within an attractor network should persist even after input is removed. To examine the temporal stability of PCx odor representations, we trained and tested a linear decoder on populations responses smoothed with a 200 ms kernel from odor onset until several seconds after odor offset in responses from TeLC-infected and contralateral control hemispheres. In control PCx, odors could be accurately identified using responses at least 4 seconds after odor offset, but odor information decayed much more rapidly after offset in TeLC-infected PCx (Fig. 6, a-c). We then asked if odor representations remained stable at later time points or whether patterns of activity evolved dynamically across time^42^. To do this we trained our decoder on responses at odor offset and then tested on subsequent epochs. We could accurately classify responses long after offset in responses from control hemispheres, whereas decoding accuracy from TeLC-PCx decayed more rapidly (Fig. 6, b-d). Similarly, odor-evoked activation patterns in SLs were less stable than those in simultaneously recorded PYRs in Ntng1-Cre mice (Fig. 6, e-h). Thus, recurrent circuitry preserves odor representations in PCx after the stimulus has ended.

**Figure 6.**
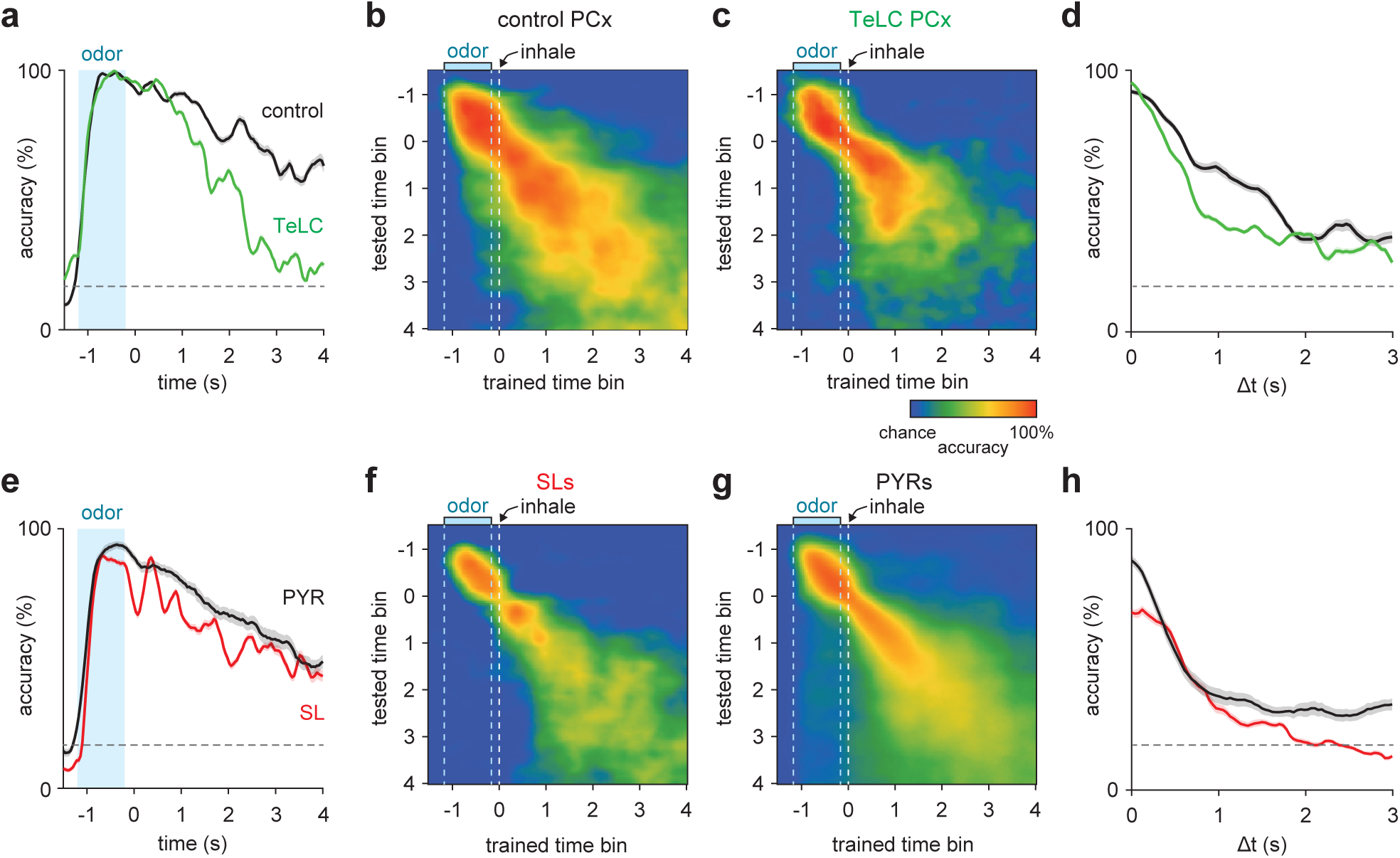
Short-term pattern stability in recurrently-connected PCx cells. (a-b) Average cross-time decoding accuracy for pseudopopulations of 200 cells recorded from control (a) or TeLC-infected (b) PCx. Responses were aligned to the first sniff after odor offset. (c) Decoding accuracy for training and testing on identical time bin at increasing times from odor offset for control (black) and TeLC- (green) PCx. Mean ± 95% bootstrapped confidence intervals. (d) Mean decoding accuracy at increasing temporal distance between the training bin (t = 0) and test bin in control (black) and TeLC- (green) PCx. Mean ± 95% bootstrapped confidence intervals. (e-h) As in a-d, but for pseudopopulations of 100 SLs (f, red) or PYRs (g, black).

## Discussion

We find that PCx is an attractor network by virtue of its recurrent activity. First, cortical sensory representations are preserved even though upstream representations are substantially degraded. Second, odor responses stabilize over multiple presentations, reflecting, in part, the formation of cortical odor templates. Third, odor representations persist after the stimulus has ended. And, two lines of evidence indicate that recurrent cortical circuitry underlies these phenomena: first, responses destabilize and do not converge in a trial-number dependent manner after eliminating recurrent circuitry; and second, odor responses are more stable in recurrently connected PYRs than in SLs.

### Stable odor representations in PCx

Recurrent connections are capable of undergoing plasticity and forming self-reinforcing neural ensembles^43-45^. This fundamental feature of cortical circuitry endows the circuit with the ability to form coherent sensory representations from noisy input^27,46^. Even under normal, stable behavioral conditions, the olfactory system is challenged with identifying complex odorant mixtures with fluctuating odorant concentrations and inherent variability at the level of receptor binding and odorant sensory neuron activity. The destabilized and degraded OB odor responses that we induced by anesthesia throw the stabilizing function of PCx processing into stark relief, but we see subtle evidence of this stabilizing function in the numerical improvement of odor separability in PCx over OB populations in the awake state (Fig. 1m). Previously, we identified complementary recurrent circuit mechanisms through which PCx transforms varying concentration-dependent OB input odor to form stable odor identity representations^37^. These results are consistent with a primary role for PCx in stable odor coding despite varying input.

It should be noted that in the present experiments animals were neither motivated to generalize odors nor make fine discriminations. This stabilization may thus serve as a default mechanism during passive odor sensing, although the balance of pattern completion or discrimination may be modulated at multiple processing stages to serve changing behavioral contexts^13,47,48^. Intriguingly, cholinergic^49^, noradrinergic^50^ and GABAergic^51,52^ modulators selectively affect recurrent but not afferent synapses in PCx, providing a mechanism to coherently and flexibly implement task-dependent computations in the same circuit.

### State-dependent odor responsivity

We saw a decrease in OB odor responsivity after injection of k/x, which afforded us the opportunity to examine how PCx responds with degraded input. However, previous work has indicated either no change^53^ or an increase in the strength of OB responses under anesthesia^32,33^. We recorded from a similar population of ventrally located M/T cells and used an identical dose of ketamine/xylazine to Rinberg et al. The most substantive methodological differences our studies are 1) their freely-moving, awake animals were able to actively truncate the odor presentation by removing their nose from the odor port, whereas in our experiments head-fixed animals passively received odors for a full second and 2) their odor responses were aligned to nose withdrawal rather than to inhalation. These would both have the effect of decreasing the apparent strength of awake responses and may largely account for the differences in our results.

It is more difficult to compare the results from our extracellular recordings with Kato et al.’s calcium imaging results. We made additional OB recordings matching the length of odor stimulus (4 s), accounting for the smoothing and loss of individual spikes associated with calcium imaging, and using a range of ketamine doses from 70-200 mg/kg and were unable to identify conditions where we could observe larger or more reliable odor responses in anesthetized OB (data not shown). We also compared the changes we observed in spontaneous and odor-evoked activity to those reported by Kollo et al. using in vivo whole-cell recordings. By contrast with these intracellular data we saw an overall decrease in firing rather than a narrowing of firing rate distributions in anesthetized OB activity. These differences may be partially explained by the large population of so-called ‘silent’ mitral cells, which we likely underrepresent with extracellular spike-sorting procedures. Nevertheless, we found that even very low firing rate cells in our recordings tend to decrease their firing under anesthesia and do not appear more likely to have strong odor responses. Future experiments will therefore be required to resolve these discrepancies. However, regardless of why our results are different, they do not impact our fundamental observation that odor representations remain robust when OB input is degraded.

### Odor representations in semilunar and pyramidal cells

Our findings indicate a functional segregation between coding properties of SLs and PYRs. Recent studies have demonstrated that local connectivity, molecular identity, and projection targets vary with depth in PCx. Superficial/SL cells receive input from OB and target PCx PYRs as well as more posterior targets such as lateral entorhinal cortex and cortical amygdala^54,55^. Because SLs receive little or no input from other PCx excitatory neurons, their responses are minimally affected by local cortical computations^41,56^. These cells may thus simply integrate converging inputs from OB with no mechanisms to correct variability in their responses inherited from OB. As such, the downstream targets of SLs receive a more variable, state-dependent olfactory representation, perhaps benefiting from specific filtering or modulation of early odor responses.

Hippocampal area CA3, has been modeled as an auto-associative network capable of storing and retrieving a large number of unique representations without interference, due to the presence of an extensive network of recurrent excitatory collaterals^5,24,25^. Similarly, because PYRs receive inputs from other local PCx excitatory neurons, they can retrieve previously stored odor-evoked activity patterns via recurrent reactivation. We propose the initial activation of a small subset of superficial PCx neurons can drive reactivation of a stable ensemble of deeper cells, enabling the recovery of robust PCx odor responses despite degraded OB input. Interestingly, PCx cells in deep layer II project back to OB^54,55,57^, suggesting that an accurate representation of the current stimulus is returned to the OB allowing comparison between ongoing input and the retrieved activation patterns, and potentially implementing a predictive loop between OB and PCx to refine odor representations in both regions^39^. This loop therefore allows SLs to receive OB input that is updated by feedback from PYRs, and may explain why odor representations in SLs are better preserved after odor offset than those in TeLC-PCx (Fig. 6)

We hypothesize that PCx circuits undergo rapid plasticity induced by stimulus exposure which embeds attractor states in the PCx synaptic architecture and later biases the trajectories of PCx population activity to previously visited states. We have demonstrated that these processes require recurrent excitation in PCx but the detailed mechanisms remain to be explored. The bulk of experience-driven changes we observe in PCx are accompanied by decreases in overall stimulus-evoked spiking, suggesting that plasticity in both excitatory and inhibitory connectivity may effect response stabilization^58,59^. Also, it remains unclear whether the instability in TeLC-PCx is due to the inability to retrieve patterns or to store them in the first place. Addressing these questions will require development of temporally-restricted and synapse-type specific interventions. Nevertheless, these results demonstrate that computations carried out by recurrently-connected cells within PCx enable pattern stabilization and strongly suggest that PCx serves as a locus of memory storage for odor representations acquired through incidental sensory experience.

## Acknowledgements

We thank J. Beck and G. Field for helpful discussions and A. Fleischmann, L. Glickfeld, S. Lisberger and A. Schaefer for helpful comments on earlier versions of the manuscript. This work was supported by grants from NIDCD (DC015525 and DC016782) and the Edward Mallinckrodt Jr. Foundation.

## Methods

All experimental protocols were approved by Duke University Institutional Animal Care and Use Committee. The methods for head-fixation, data acquisition, electrode placement, stimulus delivery, and analysis of single-unit and population odor responses are adapted from those described in detail previously ^23^. A portion of the data reported here (5 of 12 simultaneous OB-PCx experiments) were also described in that previous report.

### Mice

All mice except Ntng1-cre mice were adult (>P60, 20-24 g) offspring of Emx1-cre (+/+) breeding pairs obtained from The Jackson Laboratory (005628).

Ntng1-Cre knock-in mouse was generated using CRISPR based method in which sequences encoding the Cre recombinase followed by the bovine growth hormone polyA sequences were inserted at the start codon ATG of Ntng1 gene. During CRISPR mediated homologous recombination, an extra 162 base pairs was also inserted in front of the ATG start codon of Cre (note that these 162 base pairs are in-frame with Cre). The 162 extra DNA sequences are: atgtatttgtcaagattcctgtcgatccatgccctgtgggtgacagtgtcctctgtgatgcagccctaccttacattatcagatctgaattcactagtcgcgcccggggagcccaaaggttaccccagttggggcgggcccgaacgaaaaggtagggctgcc.

Thus, the resulting allele contains Cre with an extra 5’ end 34 amino acids inserted into the start codon of the Ntng1 gene. The knock-in allele was verified using both genomic PCR and Southern blot.

### Head-fixation

Mice were habituated to head-fixation and tube restraint for 15-30 minutes on each of the two days prior to experiments. The head post was held in place by two clamps attached to ThorLabs posts. A hinged 50 ml Falcon tube on top of a heating pad (FHC) supported and restrained the body in the head-fixed apparatus.

### Ketamine/xylazine anesthesia

To induce anesthesia, animals were injected with a bolus ketamine/xylazine cocktail (100/10 mg/kg, ip). This induced stable anesthesia lasting 30-45 minutes. Throughout the recording, body temperature was maintained using a heating pad (FHC). During anesthesia breathing became metronomic and animals ceased spontaneous forelimb movements.

We defined a subset of 7 awake trials and 7 k/x trials for all analyses except the analysis of odor representation stabilization in Fig. 5A-E. Awake trials were selected as the last 7 trials prior to k/x injection, to minimize sniff-related variability which was prevalent in early trials. K/X trials were defined as the 7 trials following behavioral and electrophysiological onset of anesthesia effects, most prominently expressed in the loss of variable breathing frequency and increase in low-frequency LFP power.

### Data acquisition

Electrophysiological signals were acquired with 32-site polytrode acute probes (A1×32-Poly3-5mm-25s-177, Neuronexus) through an A32-OM32 adaptor (Neuronexus) connected to a Cereplex digital headstage (Blackrock Microsystems). Unfiltered signals were digitized at 30 kHz at the headstage and recorded by a Cerebus multichannel data acquisition system (BlackRock Microsystems). Experimental events and respiration signal were acquired at 2 kHz by analog inputs of the Cerebus system. Respiration was monitored with a microbridge mass airflow sensor (Honeywell AWM3300V) positioned directly opposite the animal’s nose. Negative airflow corresponds to inhalation and negative changes in the voltage of the sensor output.

### Electrode placement

The recording probe was positioned in the anterior piriform cortex using a Patchstar Micromanipulator (Scientifica). For piriform cortex recordings, the probe was positioned at 1.32 mm anterior and 3.8 mm lateral from bregma. Recordings were targeted 3.5-4 mm ventral from the brain surface at this position with adjustment according to the local field potential (LFP) and spiking activity monitored online. Electrode sites on the polytrode span 275 µm along the dorsal-ventral axis. The probe was lowered until a band of intense spiking activity covering 30-40% of electrode sites near the correct ventral coordinate was observed, reflecting the densely packed layer II of piriform cortex. For simultaneous ipsilateral olfactory bulb recordings, a micromanipulator holding the recording probe was set to a 10-degree angle in the coronal plane, targeting the ventrolateral mitral cell layer. The probe was initially positioned above the center of the olfactory bulb (4.85 AP, 0.6 ML) and then lowered along this angle through the dorsal mitral cell and granule layers until encountering a dense band of high-frequency activity signifying the targeted mitral cell layer, typically between 1.5 and 2.5 mm from the bulb surface.

### Spike sorting and waveform characteristics

Individual units were isolated using Spyking-Circus (https://github.com/spyking-circus)^60^. Clusters with >1% of ISIs violating the refractory period (< 2 ms) or appearing otherwise contaminated were manually removed from the dataset. Pairs of units with similar waveforms and coordinated refractory periods in the cross-correlogram were combined into single clusters. Unit position with respect to electrode sites was characterized as the average of all electrode site positions weighted by the wave amplitude on each electrode. Relative dorsal-ventral unit position was determined by fitting the waveform positions within a recording with a truncated normal distribution (truncated at 0 and 275) and then subtracting the mean of this fit.

### Unit stability criteria

To assure stable identification of cells across states, sorted units were subjected to further stability criteria (Supplementary Fig. 2). Units were excluded if their overall rate fell below 0.01 Hz, if their rate changed more than 100-fold across states, or if their average peak-to-peak waveform amplitude changed by more than 50 μV across states. 107/294 OB units and 63/703 PCx units were discarded based on these criteria.

### Percent responding and sparseness bootstrap analyses

Cell-odor pairs were labeled as significantly odor-responsive using a rank-sum test (p < 0.05) on trial-by-trial spike counts over the first sniff after odor delivery compared to spike counts over the last pre-odor sniff. Because of variability in cell yield for olfactory bulb experiments we computed population responsivity measures on the population of all recorded cells rather than for each experiment. We established confidence bounds for these measures (percent activated responses, lifetime and population sparseness) by sampling these with replacement from the population of all recorded cells 1000 times. For significance testing, a null distribution of mean differences (between awake and k/x samples) was constructed by randomly selecting equivalent-sized samples from the combined population 1000 times, and the p-value was the fraction of null responses more extreme than the empirical mean difference.

### Trial-trial population vector correlations

The similarity of population odor responses, defined as spike count vectors within the first sniff, was quantified using the Pearson correlation coefficient. Population responses were combined across experiments to form a pseudopopulation, and correlations for all trial-pairs were calculated (i.e. 7 trials for two odors = 49 correlations). The correlation between two stimuli (across odor) or between a stimulus and itself (within odor) was then taken as the average of these correlations. Separation of population firing rate vectors in neural space was the average within-odor correlation minus the average across-odor correlation. Bootstrap significance testing on pseudo-population correlation measures were determined with the null hypothesis that mean differences between OB and PCx could be generated from a homogenous population containing cells from both regions. We combined OB and PCx responses and sampled this distribution with replacement to match the recorded population size of OB and PCx cells and then computed the mean difference between correlation separation for these populations 1000 times. The p-value was then the fraction of null differences that were more extreme than the empirical mean difference.

### Population decoding analysis

Odor classification accuracy based on population responses was measured using a linear multi-class SVM classifier with 10-fold cross-validation (LIBLINEAR, solver 4 (Crammer and Singer method), https://www.csie.ntu.edu.tw/~cjlin/liblinear/ ^61^. Responses to six distinct monomolecular odorants presented at 0.3% v/v were used as the training and testing data. The feature vectors for spike count classification were the spike counts for each cell during the 480 ms following inhalation. For decoding across states, the classifier was trained on all awake trials and accuracy was assessed across all anesthetized trials.

Classification accuracy was measured across multiple pseudopopulation sizes. To estimate the mean accuracy at each size, we constructed pseudopopulations by randomly subsampling from the entire recorded population 200 times. Bootstrap confidence intervals on the mean accuracy for each population size were estimated by sampling with replacement from the distribution of accuracies and re-computing the mean 1000 times.

### Local field potential analysis

30 kHz raw recordings were downsampled to 1 kHz and filtered between 0.05-500 Hz with a 3-pole Butterworth filter. Average power spectra and OB-PCx coherence were obtained using the multi-taper spectrum utilities in the Chronux package (www.chronux.org). For visualization, LFPs were pre-whitened with an 2nd-order autoregressive filter, and spectrograms were computed in a 30-s sliding window in 3 s steps.

### Spontaneous activity and pairwise phase consistency

Spontaneous activity was computed using spikes that occurred > 3 s after odor offset and > 2 s before odor onset. The relationship of each unit’s spiking to the ongoing respiratory oscillation was quantified using pairwise phase consistency (PPC) as in ^62^. Each spike was assigned a phase by interpolation between inhalation (0 degrees) and exhalation (180 degrees). Each spike was then treated as a unit vector and PPC was taken as the average of the dot products of all pairs of spikes.

### Demixed PCA overlap analysis

Demixed PCA projections were computed using the Machens lab, MATLAB implementation (https://github.com/machenslab/dPCA)^36^. Trial-by-trial pseudopopulation responses were constructed as above from all recorded cells using a 500 ms response window. PCs were then computed on Stimulus, State, and Stimulus X State interaction marginalizations. Trial responses were projected onto the top 3 Stimulus components and the trial-mean and covariance of these 3D projections for each odor in each state were determined. These measures were then used to compute an overlap score for each odor across states according to Matusita’s measure ^63,64^.

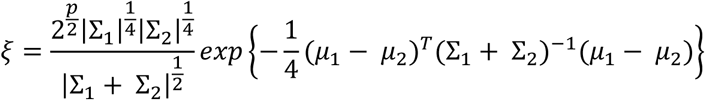

### Optogenetic tagging of NTNG1^+^ cells

In 5/12 experiments with NTNG1-Cre mice, putative semilunar cells were tagged by Cre-dependent expression of Jaws (a red-shifted variant of halorhodopsin), and in 5/12 experiments with NTNG1-Cre mice, putative semilunar cells were tagged by Cre-dependent expression of Archaeorhodopsin (Arch). Cells expressing either inhibitory opsin show rapid onset and deep suppression during exposure to 532 nm (Arch) or 640 nm (Jaws) light. Normal odor response recordings were made with an optic fiber attached probe, and 20 1 s laser pulses were delivered at the end of the experiment. Two criteria were applied to identify Arch^+^ cells: 1) p < 0.0001 in rank-sum test of spiking in the 1 s preceding and during laser stimulation, 2) median last-spike latency during laser pulse < 0.01 ms (shown as red dots). Additionally, cells with overall firing rates < 0.175 Hz or a peak-trough time < 0.35 ms in their average waveform were excluded from classification either as Arch^+^ or Arch^-^cells.

2/12 experiments used excitation with channelrhodopsin for opto-tagging. Cells were stimulated using ∼200, 1 ms pulses of 473 nm light, delivered at 4 Hz at the end of the experiment. Two criteria were applied to identify ChR2^+^ cells: 1) p-value < 0.001 in Stimulus-Associated spike Latency Test ^65^, 2) latency-to-peak response in PSTH < 0.003 ms. Cells with a peak-trough time < 0.35 in their average waveform were excluded from classification as ChR2^+^ or ChR2^-^.

### Neural distance-to-stable and regression analysis

For each experiment, population firing rate vectors for each odor trial were constructed from spiking during the 500 ms after odor inhalation or from a control period 2000 to 1500 ms before odor inhalation. Mean Euclidean distance from each trial population response to the last five awake odor responses (stable trials) was normalized to the mean Euclidean distance between stable trials to estimate population distance-to-stable. For the main analysis of PCx and OB population trajectories, PCx data was combined from simultaneous OB-PCx recordings and Ntng-Cre recordings, and three OB-PCx experiments which had <15 awake trials prior to k/x injection were omitted (n = 22 PCx recordings; n = 9 OB recordings).

To examine the influence of sniffing, overall firing rates, and odor experience over trials on the stabilization of population responses, we fit a multiple linear regression model using the *fitlm* function in the MATLAB Statistics and Machine Learning toolbox with sniff rate, firing rate, and trials, as predictors of distance-to-stable. Sniff rate was estimated as the reciprocal of the first breath duration following odor presentation. Population firing rates were taken as the mean response across neurons on each trial. All predictors were z-scored to allow comparison of the magnitude of regression coefficients. To further visualize stabilization across trials that is independent of sniffing and overall firing rates, we fit a reduced model including only sniff rate and firing rate as predictors, and examined the residuals of this model as a function of trials.

### Cross-time decoding and temporal stability

To assess stability of odor representations across short timescales, we trained and tested an SVM classifier (as above) on different time bins following odor offset. Smooth pseudopopulation responses were built from kernel density functions (200 ms kernel) aligned to the first inhalation after odor offset, and the classifier was trained and tested on each combination of time points up to 1.5 s before and 4 s after inhalation. Classification accuracy was assessed with leave-one-out cross-validation and no time bins from the test trial were included in the training data for each fold.

### Data availability

Raw data and code will be made available upon reasonable request.

## Supplementary Information

**Supplementary Figure 1.**
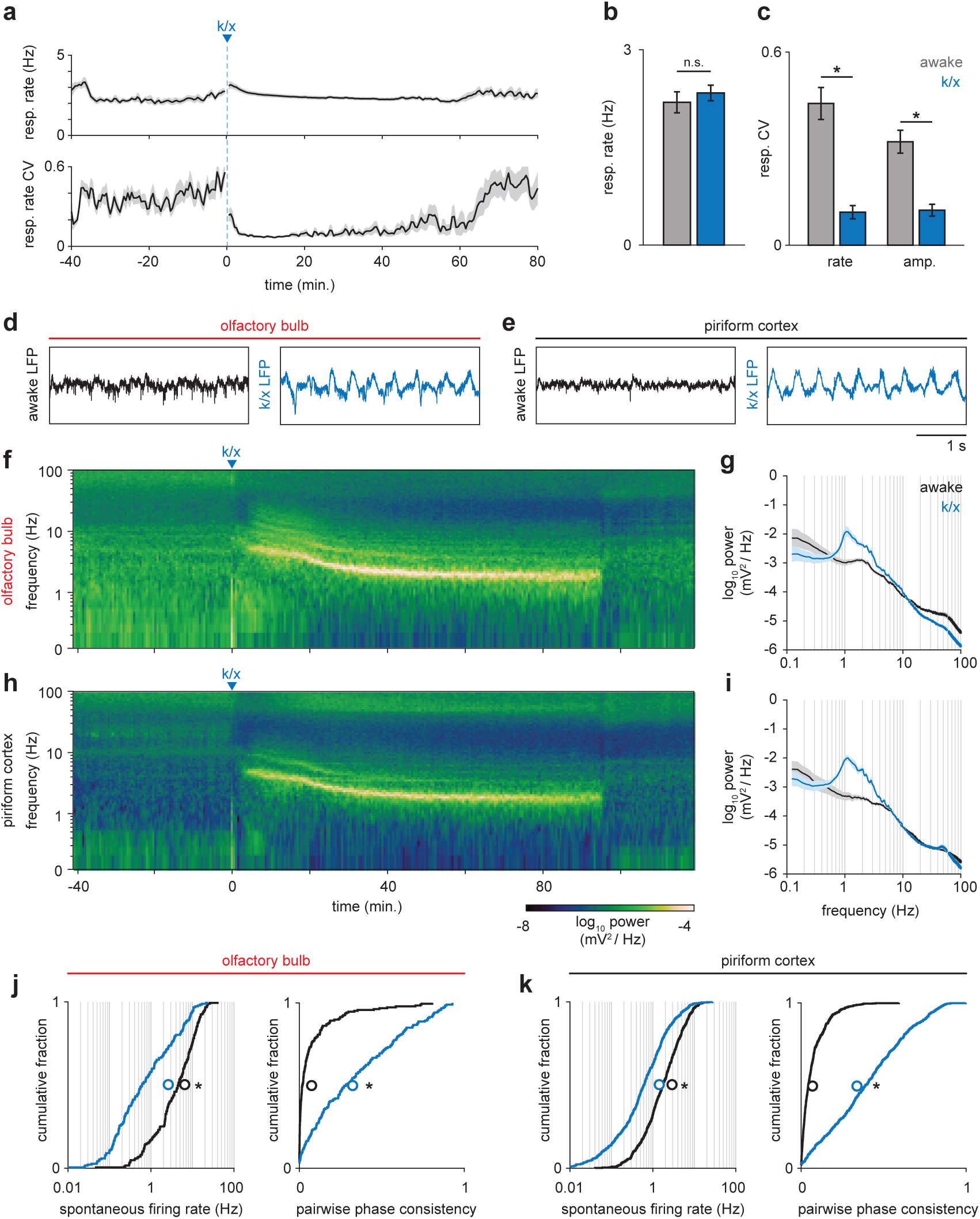
State-dependent changes in respiration, local field potential and spontaneous spiking. (a) Respiration rate (top) and coefficient of variation (CV, bottom) measured in a sliding 60 second window before and during anesthesia (n = 12 experiments, mean ± sem). (b) Average respiration rate did not differ between awake (gray) and anesthetized periods (blue); paired t-test, t(11) = −1.07, p = 0.30. (c) Variability in the rate and amplitude or respiration decreases under anesthesia (paired t-tests; rate: t(11) = 6.93, p = 2.48e-5; amplitude: t(11) = 6.98, p = 2.32e-5.) Error bars are mean ± sem. (d-e) Example local field potential traces from OB (d) and PCx (e) from awake and anesthetized periods. Awake LFP in both regions is desynchronized, while anesthetized LFP becomes strongly coupled to respiration. (f) Example spectrogram showing LFP power in OB throughout an experiment. After k/x injection, high frequency power rapidly diminishes and is replaced by strong low frequency oscillations. (g) Average power spectra during awake and anesthetized periods in OB (n = 12, mean ± sem). (h-i) As in f and g but for PCx (n = 11, mean ± sem). (j) Spontaneous spiking activity is reduced and more couple to respiration in OB under anesthesia. Left, cumulative distribution of spontaneous firing rates in awake (black) and anesthetized (blue) periods. Mean shown as unfilled circles. Paired t-test, t(186) = 14.46, p = 6.22e-34. Right, cumulative distribution of pairwise phase-consistency of spiking relative to respiration phase. Paired t-test, t(186) = −15.03, p = 3.07e-32. (k) As in j, but for PCx. Paired t-tests: Spontaneous rate, t(639) = 21.66, p = 1.88e-78. Pairwise phase consistency, t(639) = −29.52, p = 1.81e-121.

**Supplementary Figure 2.**
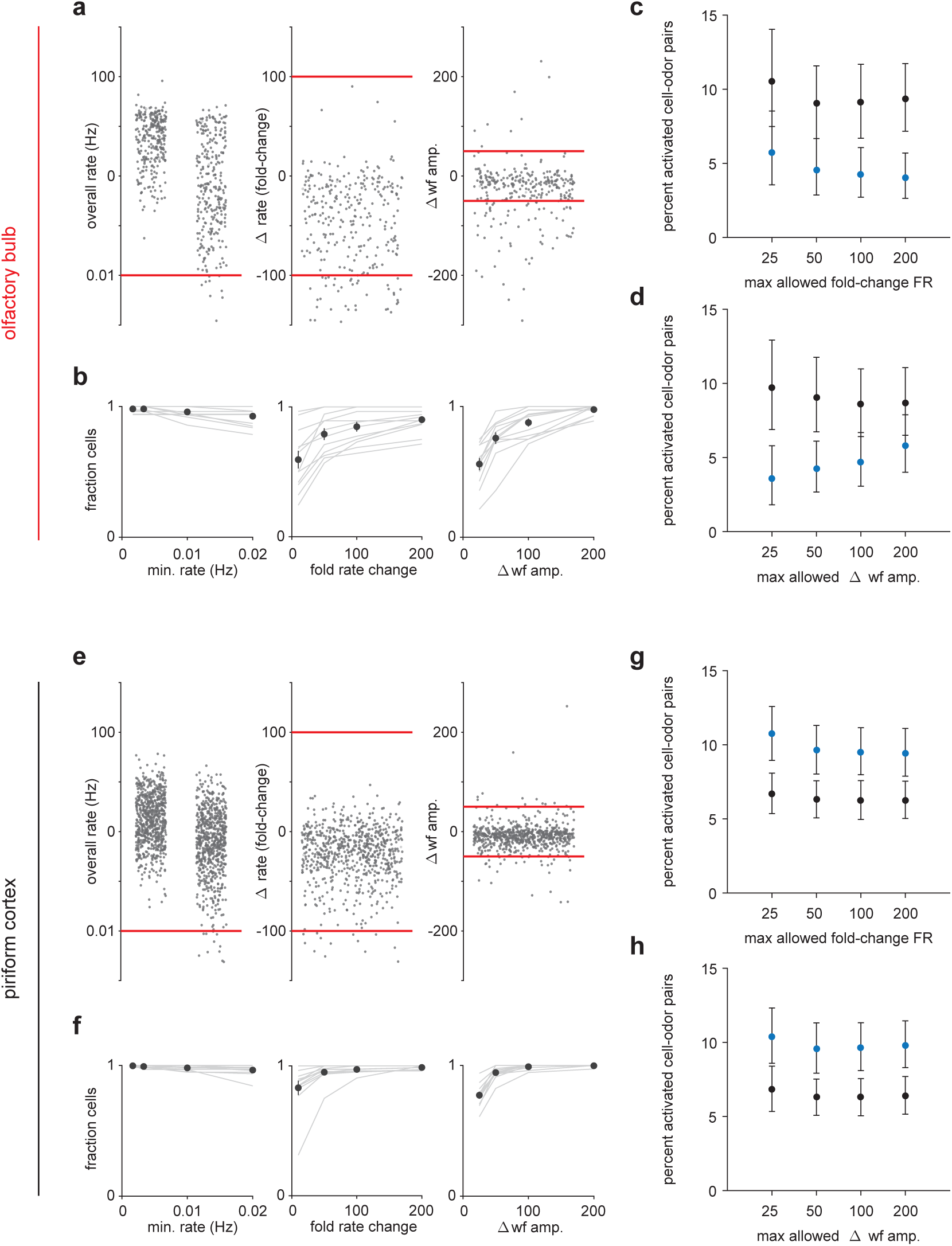
Criteria for maintaining sorted unit identity across states. (a) Left, Overall firing rates for all sorted OB units. Units falling below one spike in 0.01 Hz in either state were excluded. Middle, Fold-change in overall firing rate across states. Units with a greater than 100-fold change were excluded. Right, Change in waveform amplitude across states. Spike amplitudes for each unit were measured on the channel with the largest peak-to-peak waveforms. Units with greater than 50 μV changes in peak-to-peak amplitude were excluded. (b) Fraction of cells considered stable under varying rate (left), rate-change (middle), and waveform amplitude (right) criteria. (c-d) Primary experimental results do not depend strongly on the choice of stability criteria. The percentage of cell-odor pairs significantly activated by odor in OB is always substantially lower under anesthesia than during awake trials despite varying the rate-change (c) or waveform (d) criteria. Stricter waveform stability criteria tend to enhance the effect. (e-h) As in (a-d) but for PCx.

**Supplementary Figure 3.**
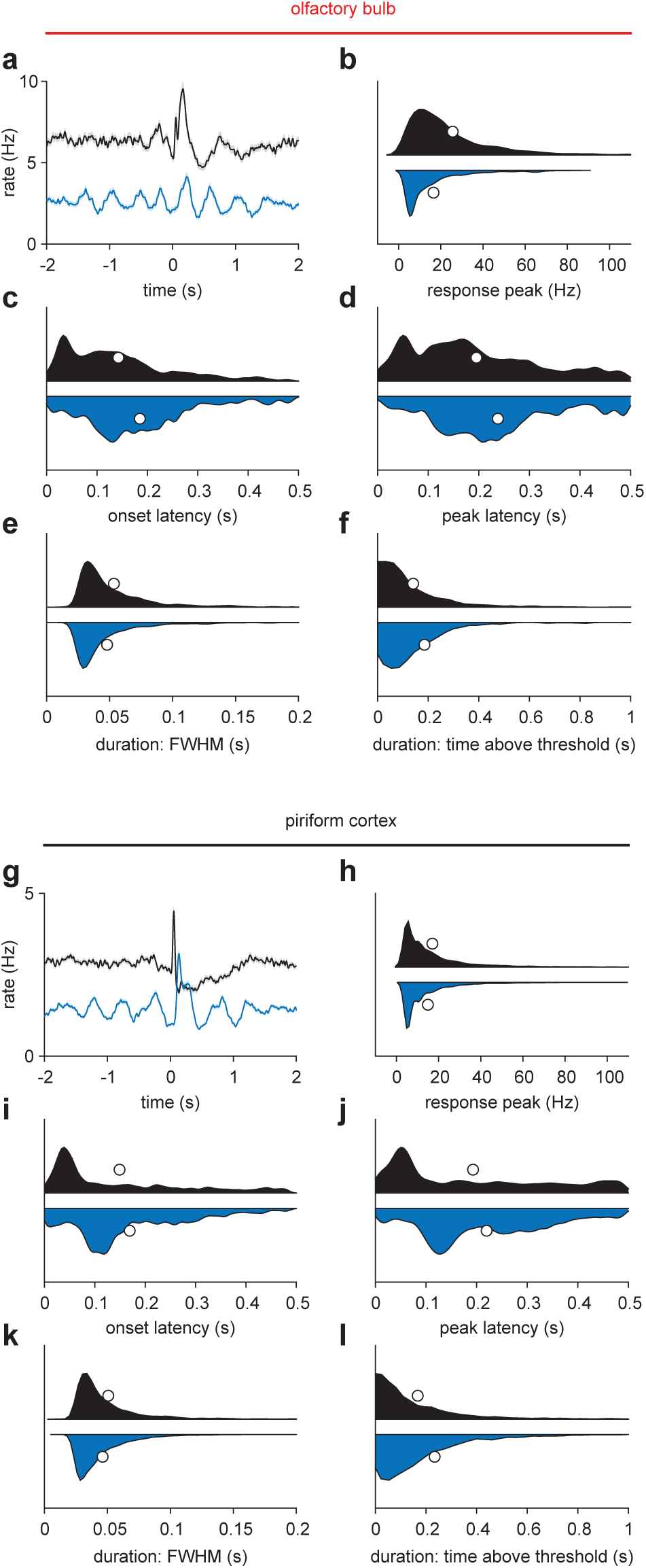
Odor response characteristics during awake and anesthetized trials. (a) Average odor response for all OB cell-odor pairs during awake (black) or k/x (blue) trials (n = 1122 cell-odor pairs, mean ± sem). (b-f) Responses with any peak in the 500 ms following inhalation were identified and their response features characterized. Awake, n = 1059; K/X, n = 799 cell-odor pairs. (b) Response peak is the maximum of the cell-odor PSTH within 500 ms. (c) Onset latency is the latency to cross a threshold of 2.5 s.d. above the 3 second pre-odor baseline. (d) Peak latency is the time of the maximum of the cell-odor PSTH within 500 ms. (e) Response duration measured as full-width at half-maximum in the first 500 ms. (f) Response duration measures as time above a 2.5 s.d. threshold. This accounts for multiphasic responses that may have sharp initial peaks followed by later, lower amplitude rate increases. Awake responses tended to be larger and lower latency, with slightly wider peaks but shorter overall responses. (g-l) As in a-f, but for PCx.

**Supplementary Figure 4.**
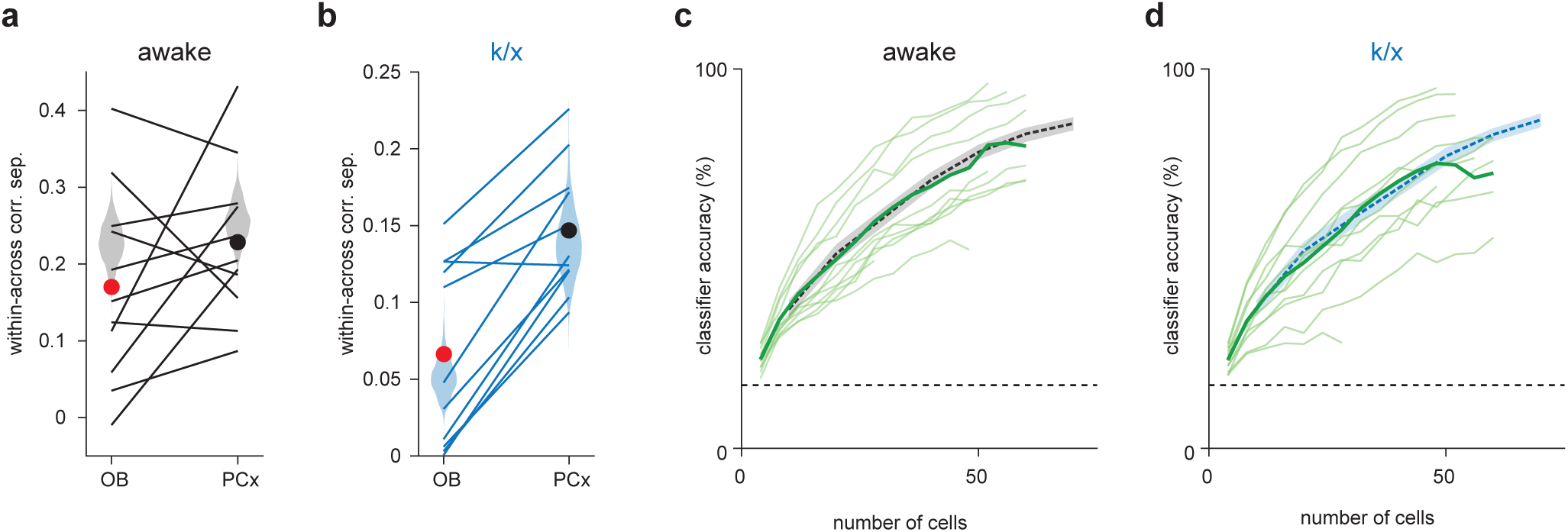
Similar outcomes for simultaneously recorded PCx populations and pseudopopulations. (a-b) Within-across odor population vector correlation difference for awake trials (a) or k/x trials (b) in OB and PCx. Each line is a separate simultaneous OB-PCx recording (n = 11). Circles show means for OB (red) and PCx (black). Pseudopopulation correlations measures from Figure 1 are shown as shaded violin plots for comparison. (c-d) Classifier accuracy for simultaneously recorded PCx populations up to 60 cells in awake (c) or k/x trials (d). Each light green line is a separate PCx recording. The average accuracy across recordings is shown as a bold green line. Classifier performance from Figure 1 is shown as a dashed line for comparison.

**Supplementary Figure 5.**
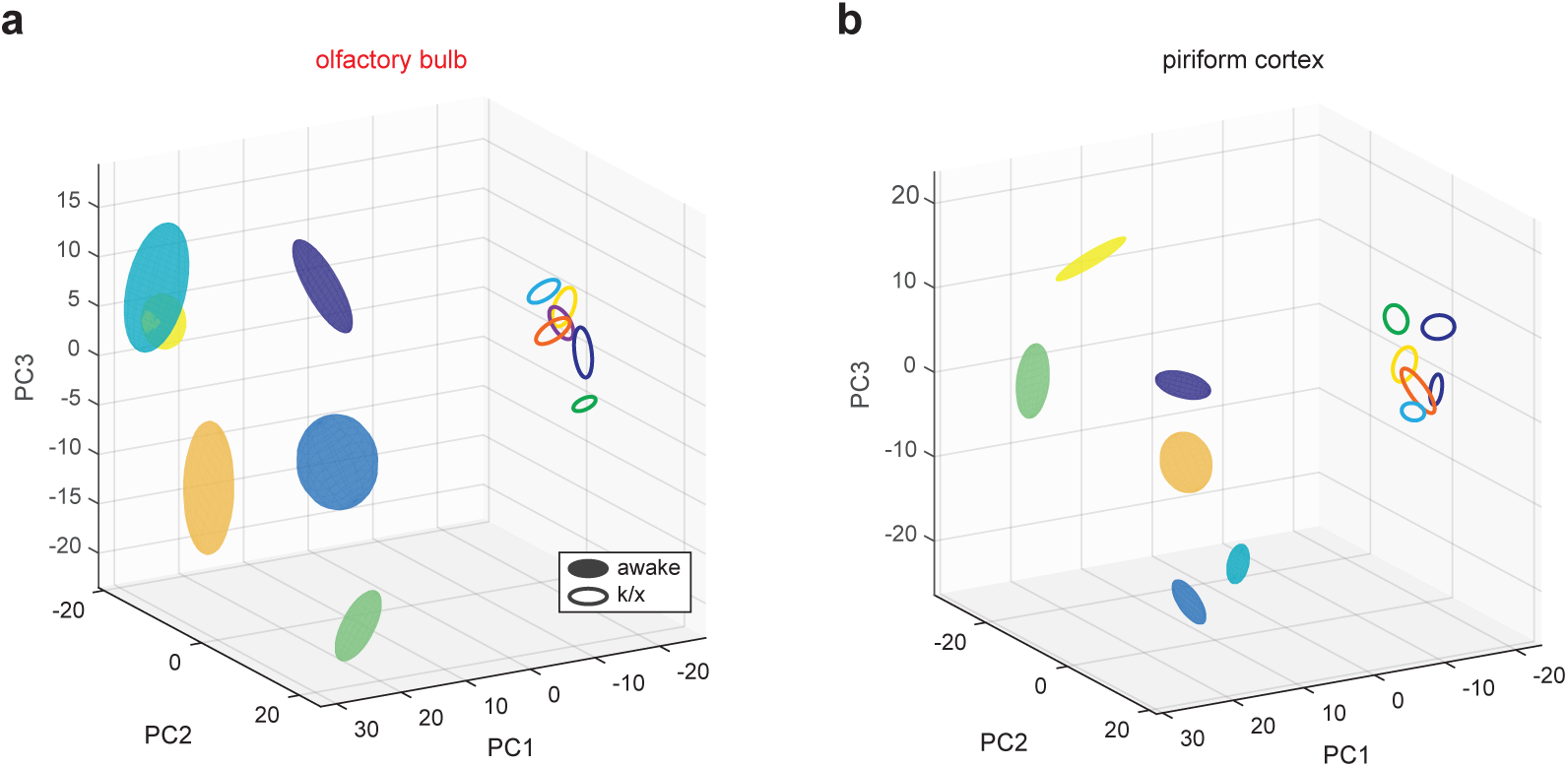
State-dependent low-dimensional representations in OB and PCx. (a) OB pseudopopulation responses to six different odors (different colors) recorded in awake filled spheres) and anesthetized states (open spheres) projected onto the first three principal components. Spheres are centered on response mean and describe ± 1 s.d. ellipsoids. (b) As in a, but for PCx.

**Supplementary Figure 6.**
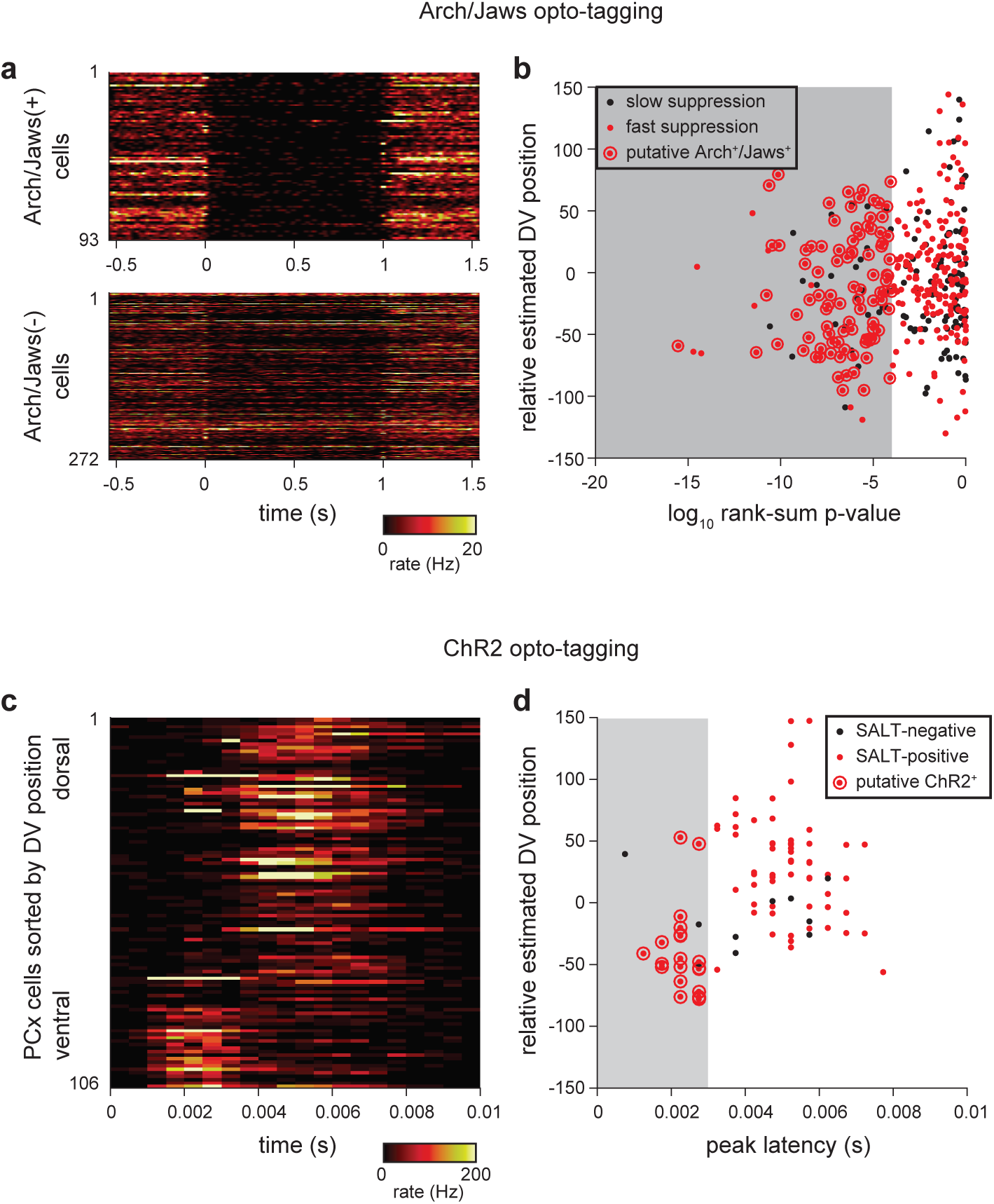
Criteria for identifying opto-tagged Ntng1+ cells. (a) Heatmaps showing trial-averaged response to 20, 1 s laser pulses in presumptive Arch+/Jaws+ (top) and Arch-/Jaws- (bottom). Arch or Jaws-expressing cells show rapid onset and deep suppression during exposure to 532 nm or 640 nm light, respectively. (b) Two criteria were applied to identify Arch+/Jaws+ cells: 1) p < 0.0001 in rank-sum test of spiking in the 1 s preceding and during laser stimulation, 2) median last-spike latency during laser pulse < 0.01 ms (shown as red dots). Additionally, cells with overall firing rates < 0.175 Hz or a peak-trough time < 0.35 ms in their average waveform were excluded from classification either as Arch+/Jaws+ or Arch-/Jaws-cells. Arch+/Jaws+ cells (red dots circled in red) identified with these criteria were subtly shifted toward more superficial recording locations compared to the total recorded population. (c) Two experiments (out of 12) used excitation with ChR2 for opto-tagging. Heatmaps show trial-averaged responses for all recorded cells to ∼200, 1 ms pulses of 473 nm light, delivered at 4 Hz. Cells are sorted within heatmap by estimated DV position. ChR2+ cells show rapid and reliable stimulus-locked spiking. (d) Two criteria were applied to identify ChR2+ cells: 1) p-value < 0.001 in Stimulus-Associated spike Latency Test (see Methods), 2) latency-to-peak response in PSTH < 0.003 ms. Cells with a peak-trough time < 0.35 in their average waveform were excluded from classification as ChR2+ or ChR2-. ChR2+ cells (red dots circled in red) tended toward more superficial recording locations compared to the total recorded population.

**Supplementary Figure 7.**
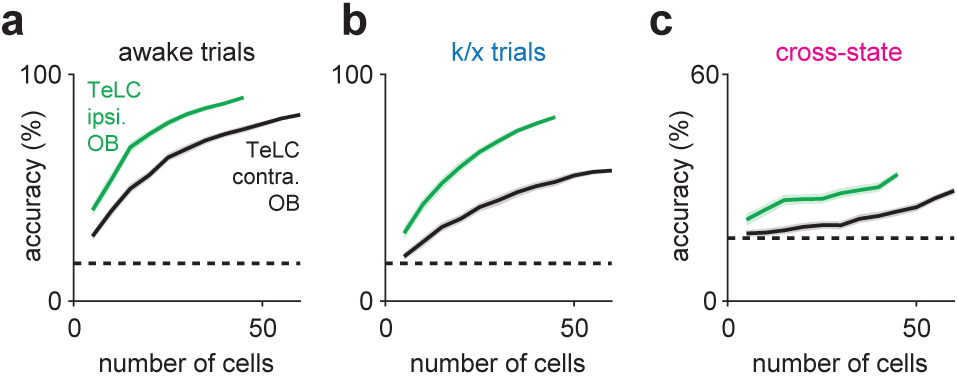
Decoding from TeLC-ipsilateral and contralateral OB populations. (a-b) Odor classification accuracy as a function of pseudopopulation size in OB ipsilateral (green) or contralateral (black) to TeLC-infected PCx in awake (a) and anesthetized (b) states. Mean ± 95% bootstrapped confidence intervals. (c) Cross-state decoding accuracy in TeLC-ipsilateral (green) or contralateral (black) OB. Mean ± 95% bootstrapped confidence intervals.

## References

1 Hopfield, J. J. Neural networks and physical systems with emergent collective computational abilities. Proc Natl Acad Sci U S A 79, 2554–2558 (1982).

2 Marr, D. Simple memory: a theory for archicortex. Philosophical transactions of the Royal Society of London. Series B, Biological sciences 262, 23–81 (1971).

3 Haberly, L. B. & Bower, J. M. Olfactory cortex: model circuit for study of associative memory? Trends Neurosci 12, 258–264 (1989).

4 Hasselmo, M. E., Wilson, M. A., Anderson, B. P. & Bower, J. M. Associative memory function in piriform (olfactory) cortex: computational modeling and neuropharmacology. Cold Spring Harbor symposia on quantitative biology 55, 599–610 (1990).

5 Treves, A. & Rolls, E. T. Computational analysis of the role of the hippocampus in memory. Hippocampus 4, 374–391, doi:10.1002/hipo.450040319 (1994).

6 Eichenbaum, H. Hippocampus: cognitive processes and neural representations that underlie declarative memory. Neuron 44, 109–120, doi:10.1016/j.neuron.2004.08.028 (2004).

7 Rolls, E. T. The mechanisms for pattern completion and pattern separation in the hippocampus. Front Syst Neurosci 7, 74, doi:10.3389/fnsys.2013.00074 (2013).

8 Guzman, S. J., Schlogl, A., Frotscher, M. & Jonas, P. Synaptic mechanisms of pattern completion in the hippocampal CA3 network. Science 353, 1117–1123, doi:10.1126/science.aaf1836 (2016).

9 Chaudhuri, R. & Fiete, I. Computational principles of memory. Nat Neurosci 19, 394–403, doi:10.1038/nn.4237 (2016).

10 Nakazawa, K. et al. Requirement for hippocampal CA3 NMDA receptors in associative memory recall. Science 297, 211–218, doi:10.1126/science.1071795 (2002).

11 Wills, T. J., Lever, C., Cacucci, F., Burgess, N. & O’Keefe, J. Attractor dynamics in the hippocampal representation of the local environment. Science 308, 873–876, doi:10.1126/science.1108905 (2005).

12 Neunuebel, J. P. & Knierim, J. J. CA3 retrieves coherent representations from degraded input: direct evidence for CA3 pattern completion and dentate gyrus pattern separation. Neuron 81, 416–427, doi:10.1016/j.neuron.2013.11.017 (2014).

13 Chapuis, J. & Wilson, D. A. Bidirectional plasticity of cortical pattern recognition and behavioral sensory acuity. Nat Neurosci 15, 155–161, doi:10.1038/nn.2966 (2012).

14 Suzuki, M. & Gottlieb, J. Distinct neural mechanisms of distractor suppression in the frontal and parietal lobe. Nat Neurosci 16, 98–104, doi:10.1038/nn.3282 (2013).

15 Carrillo-Reid, L., Yang, W., Bando, Y., Peterka, D. S. & Yuste, R. Imprinting and recalling cortical ensembles. Science 353, 691–694, doi:10.1126/science.aaf7560 (2016).

16 Inagaki, H. K., Fontolan, L., Romani, S. & Svoboda, K. Discrete attractor dynamics underlies persistent activity in the frontal cortex. Nature 566, 212–217, doi:10.1038/s41586-019-0919-7 (2019).

17 Sosulski, D. L., Bloom, M. L., Cutforth, T., Axel, R. & Datta, S. R. Distinct representations of olfactory information in different cortical centres. Nature 472, 213–216, doi:10.1038/nature09868 (2011).

18 Ghosh, S. et al. Sensory maps in the olfactory cortex defined by long-range viral tracing of single neurons. Nature 472, 217–220, doi:10.1038/nature09945 (2011).

19 Miyamichi, K. et al. Cortical representations of olfactory input by trans-synaptic tracing. Nature 472, 191–196, doi:10.1038/nature09714 (2011).

20 Stettler, D. D. & Axel, R. Representations of Odor in the Piriform Cortex. Neuron 63, 854–864 (2009).

21 Roland, B., Deneux, T., Franks, K. M., Bathellier, B. & Fleischmann, A. Odor identity coding by distributed ensembles of neurons in the mouse olfactory cortex. eLife 6, doi:10.7554/eLife.26337 (2017).

22 Miura, K., Mainen, Z. F. & Uchida, N. Odor representations in olfactory cortex: distributed rate coding and decorrelated population activity. Neuron 74, 1087–1098, doi:10.1016/j.neuron.2012.04.021 (2012).

23 Bolding, K. A. & Franks, K. M. Complementary codes for odor identity and intensity in olfactory cortex. eLife 6, doi:10.7554/eLife.22630 (2017).

24 Johnson, D. M. G., Illig, K. R., Behan, M. & Haberly, L. B. New features of connectivity in piriform cortex visualized by intracellular injection of pyramidal cells suggest that “primary” olfactory cortex functions like “association” cortex in other sensory systems. J. Neurosci. 20, 6974–6982 (2000).

25 Franks, K. M. et al. Recurrent circuitry dynamically shapes the activation of piriform cortex. Neuron 72, 49–56, doi:S0896-6273(11)00741-0 [pii] 10.1016/j.neuron.2011.08.020 (2011).

26 Ambros-Ingerson, J., Granger, R. & Lynch, G. Simulation of paleocortex performs hierarchical clustering. Science 247, 1344–1348 (1990).

27 Haberly, L. B. Parallel-distributed processing in olfactory cortex: new insights from morphological and physiological analysis of neuronal circuitry. Chem Senses 26, 551–576 (2001).

28 Hasselmo, M. E. & Barkai, E. Cholinergic modulation of activity-dependent synaptic plasticity in the piriform cortex and associative memory function in a network biophysical simulation. J Neurosci 15, 6592–6604 (1995).

29 Barkai, E. & Hasselmo, M. E. Modulation of the input/output function of rat piriform cortex pyramidal cells. J Neurophysiol 72, 644–658, doi:10.1152/jn.1994.72.2.644 (1994).

30 Poo, C. & Isaacson, J. S. Odor representations in olfactory cortex: “sparse” coding, global inhi-bition, and oscillations. Neuron 62, 850–861, doi:10.1016/j.neuron.2009.05.022 (2009).

31 Murakami, M., Kashiwadani, H., Kirino, Y. & Mori, K. State-dependent sensory gating in olfactory cortex. Neuron 46, 285–296, doi:10.1016/j.neuron.2005.02.025 (2005).

32 Kato, H. K., Chu, M. W., Isaacson, J. S. & Komiyama, T. Dynamic sensory representations in the olfactory bulb: modulation by wakefulness and experience. Neuron 76, 962–975, doi:10.1016/j.neuron.2012.09.037 (2012).

33 Rinberg, D., Koulakov, A. & Gelperin, A. Sparse odor coding in awake behaving mice. J Neurosci 26, 8857–8865, doi:10.1523/JNEUROSCI.0884-06.2006 (2006).

34 Fontanini, A. & Bower, J. M. Variable coupling between olfactory system activity and respiration in ketamine/xylazine anesthetized rats. J Neurophysiol 93, 3573–3581, doi:10.1152/jn.01320.2004 (2005).

35 Li, A. et al. Effects of different anesthetics on oscillations in the rat olfactory bulb. J Am Assoc Lab Anim Sci 51, 458–463 (2012).

36 Kobak, D. et al. Demixed principal component analysis of neural population data. Elife 5, doi:10.7554/eLife.10989 (2016).

37 Bolding, K. A. & Franks, K. M. Recurrent cortical circuits implement concentration-invariant odor coding. Science 361, doi:10.1126/science.aat6904 (2018).

38 van Vreeswijk, C. & Sompolinsky, H. Chaos in neuronal networks with balanced excitatory and inhibitory activity. Science 274, 1724–1726, doi:10.1126/science.274.5293.1724 (1996).

39 Grabska-Barwinska, A. et al. A probabilistic approach to demixing odors. Nat Neurosci 20, 98–106, doi:10.1038/nn.4444 (2017).

40 Suzuki, N. & Bekkers, J. M. Neural coding by two classes of principal cells in the mouse piriform cortex. J Neurosci 26, 11938–11947, doi:10.1523/JNEUROSCI.3473-06.2006 (2006).

41 Choy, J. M. et al. Optogenetic Mapping of Intracortical Circuits Originating from Semilunar Cells in the Piriform Cortex. Cereb Cortex, doi:10.1093/cercor/bhv258 (2015).

42 Friedrich, R. W. & Laurent, G. Dynamic optimization of odor representations by slow temporal patterning of mitral cell activity. Science 291, 889–894, doi:10.1126/science.291.5505.889 (2001).

43 Kanter, E. D. & Haberly, L. B. NMDA-dependent induction of long-term potentiation in afferent and association fiber systems of piriform cortex in vitro. Brain Res 525, 175–179, doi:0006-8993(90)91337-G [pii] (1990).

44 Jung, M. W., Larson, J. & Lynch, G. Long-term potentiation of monosynaptic EPSPs in rat piriform cortex in vitro. Synapse 6, 279–283, doi:10.1002/syn.890060307 (1990).

45 Poo, C. & Isaacson, J. S. An early critical period for long-term plasticity and structural modification of sensory synapses in olfactory cortex. J. Neurosci. 27, 7553–7558, doi:10.1523/jneurosci.1786-07.2007 (2007).

46 Wilson, D. A. & Sullivan, R. M. Cortical processing of odor objects. Neuron 72, 506–519, doi:S0896-6273(11)00962-7 [pii] 10.1016/j.neuron.2011.10.027 (2011).

47 Koldaeva, A., Schaefer, A. T. & Fukunaga, I. Rapid task-dependent tuning of the mouse olfactory bulb. eLife 8, doi:10.7554/eLife.43558 (2019).

48 Chu, M. W., Li, W. L. & Komiyama, T. Balancing the Robustness and Efficiency of Odor Representations during Learning. Neuron 92, 174–186, doi:10.1016/j.neuron.2016.09.004 (2016).

49 Hasselmo, M. E. & Bower, J. M. Cholinergic suppression specific to intrinsic not afferent fiber synapses in rat piriform (olfactory) cortex. J Neurophysiol 67, 1222–1229, doi:10.1152/jn.1992.67.5.1222 (1992).

50 Hasselmo, M. E., Linster, C., Patil, M., Ma, D. & Cekic, M. Noradrenergic suppression of synaptic transmission may influence cortical signal-to-noise ratio. J Neurophysiol 77, 3326–3339 (1997).

51 Tang, A. C. & Hasselmo, M. E. Selective suppression of intrinsic but not afferent fiber synaptic transmission by baclofen in the piriform (olfactory) cortex. Brain Res 659, 75–81, doi:10.1016/0006-8993(94)90865-6 (1994).

52 Franks, K. M. & Isaacson, J. S. Synapse-specific downregulation of NMDA receptors by early experience: A critical period for plasticity of sensory input to olfactory cortex. Neuron 47, 101–114, doi:10.1016/j.neuron.2005.05.024 (2005).

53 Kollo, M., Schmaltz, A., Abdelhamid, M., Fukunaga, I. & Schaefer, A. T. ‘Silent’ mitral cells dominate odor responses in the olfactory bulb of awake mice. Nat Neurosci 17, 1313–1315, doi:10.1038/nn.3768 (2014).

54 Diodato, A. et al. Molecular signatures of neural connectivity in the olfactory cortex. Nature communications 7, 12238, doi:10.1038/ncomms12238 (2016).

55 Mazo, C., Grimaud, J., Shima, Y., Murthy, V. N. & Lau, C. G. Distinct projection patterns of different classes of layer 2 principal neurons in the olfactory cortex. Sci Rep 7, 8282, doi:10.1038/s41598-017-08331-0 (2017).

56 Suzuki, N. & Bekkers, J. M. Two layers of synaptic processing by principal neurons in piriform cortex. J Neurosci 31, 2156–2166, doi:31/6/2156 [pii] 10.1523/JNEUROSCI.5430-10.2011 (2011).

57 Luskin, M. B. & Price, J. L. The topographic organization of associational fibers of the olfactory system in the rat, including centrifugal fibers to the olfactory bulb. Journal of Comparative Neurology 216, 264–291 (1983).

58 Vogels, T. P., Sprekeler, H., Zenke, F., Clopath, C. & Gerstner, W. Inhibitory plasticity balances excitation and inhibition in sensory pathways and memory networks. Science 334, 1569–1573, doi:10.1126/science.1211095 (2011).

59 D’Amour J. A. & Froemke, R. C. Inhibitory and excitatory spike-timing-dependent plasticity in the auditory cortex. Neuron 86, 514–528, doi:10.1016/j.neuron.2015.03.014 (2015).

60 Yger, P. et al. A spike sorting toolbox for up to thousands of electrodes validated with ground truth recordings in vitro and in vivo. eLife 7, doi:10.7554/eLife.34518 (2018).

61 Fan, R. E., Chang, K. W., Hsieh, C. J., Wang, X. R. & Lin, C. J. LIBLINEAR: A library for large linear classification. Journal of machine learning research 9, 1871–1874 (2008).

62 Vinck, M., van Wingerden, M., Womelsdorf, T., Fries, P. & Pennartz, C. M. A. The pairwise phase consistency: A bias-free measure of rhythmic neuronal synchronization. Neuroimage 51, 112–122, doi:10.1016/j.neuroimage.2010.01.073 (2010).

63 Minami, M. & Shimizu, K. Estimation of similarity measure for multivariate normal distributions. Environ Ecol Stat 6, 229–248, doi:Doi 10.1023/A:1009678412953 (1999).

64 Matusita, K. A distance and related statistics in multivariate analysis. (Academic Press, 1966).

65 Kvitsiani, D. et al. Distinct behavioural and network correlates of two interneuron types in prefrontal cortex. Nature 498, 363–366, doi:10.1038/nature12176 (2013).

